# Direct Extraction of Signal and Noise Correlations from Two-Photon Calcium Imaging of Ensemble Neuronal Activity

**DOI:** 10.1101/2021.03.11.434932

**Authors:** Anuththara Rupasinghe, Nikolas A Francis, Ji Liu, Zac Bowen, Patrick O Kanold, Behtash Babadi

**Affiliations:** Department of Electrical & Computer Engineering, University of Maryland, College Park, MD, USA; The Institute for Systems Research, University of Maryland, College Park, MD, USA; Department of Biology, University of Maryland, College Park, MD, USA; Department of Biomedical Engineering, Johns Hopkins University, Baltimore, MD, USA

**Author notes:** **For correspondence:** (BB).

## Abstract

Neuronal activity correlations are key to understanding how populations of neurons collectively encode information. While two-photon calcium imaging has created a unique opportunity to record the activity of large populations of neurons, existing methods for inferring correlations from these data face several challenges. First, the observations of spiking activity produced by two-photon imaging are temporally blurred and noisy. Secondly, even if the spiking data were perfectly recovered via deconvolution, inferring network-level features from binary spiking data is a challenging task due to the non-linear relation of neuronal spiking to endogenous and exogenous inputs. In this work, we propose a methodology to explicitly model and directly estimate signal and noise correlations from two-photon fluorescence observations, without requiring intermediate spike deconvolution. We provide theoretical guarantees on the performance of the proposed estimator and demonstrate its utility through applications to simulated and experimentally recorded data from the mouse auditory cortex.

## Introduction

Neuronal activity correlations are essential in understanding how populations of neurons encode information. Correlations provide insights into the functional architecture and computations carried out by neuronal networks (***Abbott and Dayan, 1999***; ***Averbeck et al., 2006***; ***Cohen and Kohn, 2011***; ***Hansen et al., 2012***; ***Kohn et al., 2016***; ***Kohn and Smith, 2005***; ***Lyamzin et al., 2015***; ***Montijn et al., 2014***; ***Smith and Sommer, 2013***; ***Sompolinsky et al., 2001***; ***Yatsenko et al., 2015***). Neuronal activity correlations are often categorized in two groups: *signal* correlations and *noise* correlations (***Cohen and Kohn, 2011***; ***Cohen and Maunsell, 2009***; ***Gawne and Richmond, 1993***; ***Josić et al., 2009***; ***Lyamzin et al., 2015***; ***Vinci et al., 2016***). Given two neurons, signal correlation quantifies the similarity of neural responses that are time-locked to a repeated stimulus across trials, whereas noise correlation quantifies the stimulus-independent trial-to-trial variability shared by neural responses that are believed to arise from common latent inputs.

Two-photon calcium imaging has become increasingly popular in recent years to record *in vivo* neural activity simultaneously from hundreds of neurons (***Ahrens et al., 2013***; ***Romano et al., 2017***; ***Stosiek et al., 2003***; ***Svoboda and Yasuda, 2006***). This technology takes advantage of intracellular calcium flux mostly arising from spiking activity and captures calcium signaling in neurons in living animals using fluorescence microscopy. The observed fluorescence traces of calcium concentrations, however, are indirectly related to neuronal spiking activity. Extracting spiking activity from fluorescence traces is a challenging signal deconvolution problem, and has been the focus of active research (***Deneux et al., 2016***; ***Friedrich et al., 2017***; ***Grewe et al., 2010***; ***Jewell et al., 2020***; ***Jewell and Witten, 2018***; ***Kazemipour et al., 2018***; ***Pachitariu et al., 2018***; ***Pnevmatikakis et al., 2016***; ***Stringer and Pachitariu, 2019***; ***Theis et al., 2016***; ***Vogelstein et al., 2010,2009***).

The most commonly used approach to infer signal and noise correlations from two-photon data is to directly apply the classical definitions of correlations for firing rates (***Lyamzin et al., 2015***), to fluorescence traces (***De Vico Fallani et al., 2015***; ***Francis et al., 2018***; ***Rothschild et al., 2010***; ***Winkowski and Kanold, 2013***). However, it is well known that fluorescence observations are noisy and blurred surrogates of spiking activity, because of dependence on observation noise, calcium dynamics and the temporal properties of calcium indicators. Due to temporal blurring, the resulting signal and noise correlation estimates are highly biased. An alternative approach is to carry out the inference in a two-stage fashion: first, infer spikes using a deconvolution technique, and then compute firing rates and evaluate the correlations (***Kerlin et al., 2019**; **Najafi et al., 2020**; **Ramesh et al., 2018**; **Soudry et al., 2015**; **Yatsenko et al., 2015**).* These two-stage estimates are highly sensitive to the accuracy of spike deconvolution, and require high temporal resolution and signal-to-noise ratios (***Lütcke et al., 2013**; **Pachitariu et al., 2018**).* Furthermore, these deconvolution techniques are biased towards obtaining accurate first-order statistics (i.e., spike timings) via spatiotemporal priors, which may be detrimental to recovering second-order statistics (i.e., correlations). Finally, both approaches also undermine the non-linear dynamics of spiking activity as governed by stimuli, past activity and other latent processes (***Truccolo et al., 2005**).* There are a few existing studies that aim at improving estimation of neuronal correlations, but they either do not consider signal correlations (***Rupasinghe and Babadi, 2020**; **Yatsenko et al., 2015**),* or aim at estimating surrogates of correlations from spikes such as the connectivity/coupling matrix (***Aitchison et al., 2017**; **Mishchenko et al., 2011**; **Soudry et al., 2015**; **Keeley et al., 2020**).*

Here, we propose a methodology to *directly* estimate both signal and noise correlations from two-photon imaging observations, without requiring an intermediate step of spike deconvolution. We pose the problem under the commonly used experimental paradigm in which neuronal activity is recorded during trials of a repeated stimulus. We avoid the need to perform spike deconvolution by integrating techniques from point processes and state-space modeling that explicitly relate the signal and noise correlations to the observed fluorescence traces in a multi-tier model. Thus, we cast signal and noise correlations within a parameter estimation setting. To solve the resulting estimation problem in an efficient fashion, we develop a solution method based on variational inference ***Jordan et al., 1999***; ***Blei et al., 2017***), by combining techniques from Pólya-Gamma augmentation (***Polson et al., 2013***) and compressible state-space estimation (***Rauch et al., 1965***; ***Kazemipour et al., 2018***; ***Ba et al., 2014***). We also provide theoretical guarantees on the bias and variance performance of the resulting estimator.

We demonstrate the utility of our proposed estimation framework through application to simulated and real data from the mouse auditory cortex during presentations of tones and acoustic noise. Our results corroborate existing hypotheses regarding the invariance of the noise correlation structure under spontaneous activity and stimulus-driven conditions, and its distinction from the signal correlation structure in the stimulus-driven condition (***Keeley et al., 2020***; ***Rumyantsev et al., 2020***; ***Bartolo et al., 2020***). Furthermore, while application of our proposed method to spatial analysis of signal and noise correlations in the mouse auditory cortex is consistent with existing work (***Winkowski and Kanold, 2013***), it reveals novel and distinct spatial trends in the correlation structure of layers 2/3 and 4. In summary, our method improves on existing work by: 1) joint estimation of signal and noise correlations directly from two-photon fluorescence observations without requiring intermediate spike deconvolution, 2) providing theoretical guarantees on the performance of the proposed estimator, and 3) gaining access to closed-form posterior approximations, with low-complexity and iterative update rules and minimal dependence on training data. Our proposed method can thus be used as a robust and scalable alternative to existing approaches for extracting signal and noise correlations from two-photon imaging data.

## Results

In this section we first demonstrate the utility of our proposed estimation framework through simulation studies as well as applications on experimentally-recorded data from the mouse auditory cortex. Then, we present theoretical performance bounds on the proposed estimator. Before presenting the results, we will give an overview of the proposed signal and noise correlation inference framework, and outline our contributions and their relationship to existing work. For the ease of reproducibility, we have archived a MATLAB implementation of our proposed method in GitHub (***Rupasinghe, 2020***), and have deposited the data used in this work in the Digital Repository at the University of Maryland (***Rupasinghe et al., 2021***).

### Signal and Noise correlations

We consider a canonical experimental setting in which the same external stimulus, denoted by **s**_*t*_, is repeatedly presented across *L* independent trials and the spiking activity of a population of *N* neurons are indirectly measured using two-photon calcium fluorescence imaging. ***Figure 1*** (forward arrow) shows the generative model that is used to quantify this procedure. The fluorescence observation in the *l*^th^ trial from the *j^th^* neuron at time frame *t*, denoted by 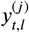, is a noisy surrogate of the intracellular calcium concentrations. The calcium concentrations in turn are temporally blurred surrogates of the underlying spiking activity 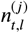, as shown in ***Figure 1***.

**Figure 1.**
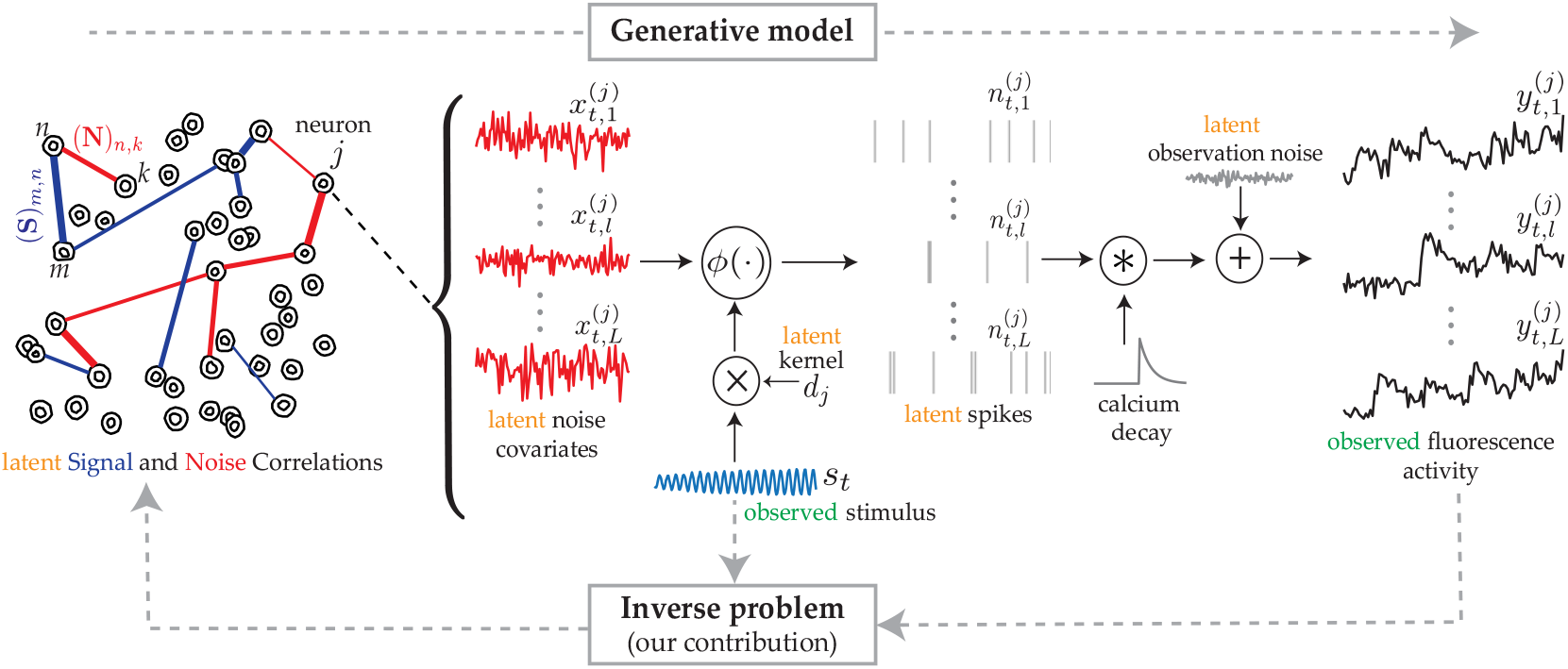
The proposed generative model and inverse problem. Observed (green) and latent (orange) variables pertinent to the *j*^th^ neuron are indicated, according to the proposed model for estimating the signal (blue) and noise (red) correlations from two-photon calcium fluorescence observations. Calcium fluorescence traces 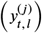 of *L* trials are observed, in which the repeated external stimulus (**s**_*t*_) is known. The underlying spiking activity 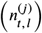, trial-to-trial variability and other intrinsic/extrinsic neural covariates that are not time-locked with the external stimulus 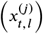, and the stimulus kernel (**d**_*j*_) are latent. Our main contribution is to solve the inverse problem: recovering the underlying latent signal (**S**) and noise (**N**) correlations directly from the fluorescence observations, without requiring intermediate spike deconvolution.

In modeling the spiking activity, we consider two main contributions: 1) the common known stimulus **s**_*t*_ affects the activity of the *j*^th^ neuron via an unknown kernel ***d***_*j*_, akin to the receptive field; 2) the trial-to-trial variability and other intrinsic/extrinsic neural covariates that are not time-locked to the stimulus **s**_*t*_ are captured by a trial-dependent latent process 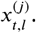 Then, we use a Generalized Linear Model to link these underlying neural covariates to spiking activity (***Truccolo et al., 2005***). More specifically, we model spiking activity as a Bernoulli process:

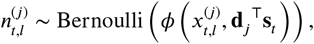

where *ϕ*(*·*) is a mapping function, which could in general be non-linear.

The *signal* correlations aim to measure the correlations in the temporal response that is time-locked to the repeated stimulus, **s**_*t*_. On the other hand, *noise* correlations in our setting quantify connectivity arising from covariates that are unrelated to the stimulus, including the trial-to-trial variability (***Keeley et al., 2020***). Based on the foregoing model, we propose to formulate the signal ((**∑**_*s*_)_i,j_) and noise covariance between ((**∑**_*s*_)_i,j_) the *i*^th^ neuron and *j*^th^ neuron as:

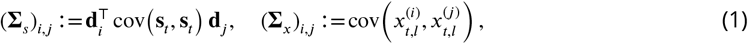

where cov(·) is the empirical covariance function defined as cov 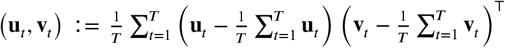, for a total observation duration of *T* time frames.

Our main contribution is to provide an efficient solution for the so-called inverse problem: direct estimation of **∑**_*s*_ and **∑**_*x*_ from the fluorescence observations, without requiring intermediate spike deconvolution (***Figure 1***, backward arrow). The signal and noise correlation matrices, denoted by **S** and **N**, can then be obtained by standard normalization of **∑**_*s*_ and **∑**_*x*_:

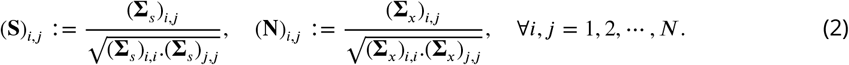

We note that when spiking activity is directly observed using electrophysiology recordings, the conventional signal 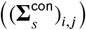 and noise 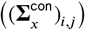 covariances of spiking activity between the *i*^th^ and *j*^th^ neuron are defined as (***Lyamzin et al., 2015***):

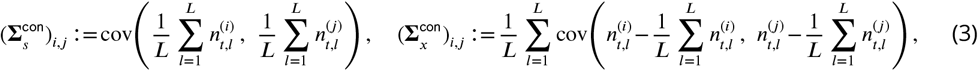

which after standard normalization in ***Equation 2*** give the conventional signal ((**S**^con^)_*i,j*_) and noise ((**N**^con^)_i,j_) correlations. While at first glance our definitions of signal and noise covariances in ***Equation 1*** seem to be a far departure from the conventional ones in ***Equation 3***, we show that the conventional notions of correlation indeed approximate the same quantities as in our definitions:

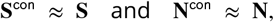

under asymptotic conditions (i.e., *T* and *L* sufficiently large). We prove this assertion of asymptotic equivalence in ***Appendix 1***, which highlights another facet of our contributions: our proposed estimators are designed to robustly operate in the regime of finite (and typically small) *T* and *L*, aiming for the very same quantities that the conventional estimators could only recover accurately under ideal asymptotic conditions.

### Existing methods used for performance comparison

In order to compare the performance of our proposed method with existing work, we consider three widely available methods for extracting neuronal correlations. In simulation studies, we additionally benchmark these estimates with respect to the known ground truth. The existing methods considered are the following:

#### Pearson Correlations from the Two-Photon Data

In this method, fluorescence observations are assumed to be the direct measurements of spiking activity, and thus empirical Pearson correlations of the two-photon data are used to compute the signal and noise correlations (***Rothschild et al., 2010***; ***Winkowski and Kanold, 2013***; ***Francis et al., 2018***; ***Bowen et al., 2020***). Explicitly, these estimates are obtained by simply replacing 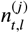 in ***Equation 3*** by 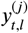, without performing spike deconvolution.

#### Two-stage Pearson Estimation

Unlike the previous method, in this case spikes are first inferred using a deconvolution technique. Then, following temporal smoothing via a narrow Gaussian kernel the Pearson correlations are computed using the conventional definitions of ***Equation 3***. For spike deconvolution, we primarily used the FCSS algorithm (***Kazemipour et al., 2018***). In order to also demonstrate the sensitivity of these estimates to the deconvolution technique that is used, we provide a comparison with the f-oopsi deconvolution algorithm (***Pnevmatikakis et al., 2016***) in ***Figure 2-Figure Supplement 1***.

#### Two-stage GPFA Estimation

Similar to the previous method, spikes are first inferred using a deconvolution technique. Then, a latent variable model called Gaussian Process Factor Analysis (GPFA) (***Yu et al., 2009***) is applied to the inferred spikes in order to estimate the latent covariates and receptive fields. Based on those estimates, the signal and residual noise correlations are derived through a formulation similar to ***Equation 1*** and ***Equation 2*** (***Ecker et al., 2014***).

### Simulation study 1: Neuronal ensemble driven by external stimulus

We simulated calcium fluorescence observations according to the proposed generative model given in Methods and Materials, from an ensemble of *N* = 8 neurons for a duration of *T* = 5000 time frames. We considered *L* = 20 repeated trials driven by the same external stimulus, which we modeled by an autoregressive process (see Methods and Materials for details). ***Figure 2*** shows the corresponding estimation results.

**Figure 2.**
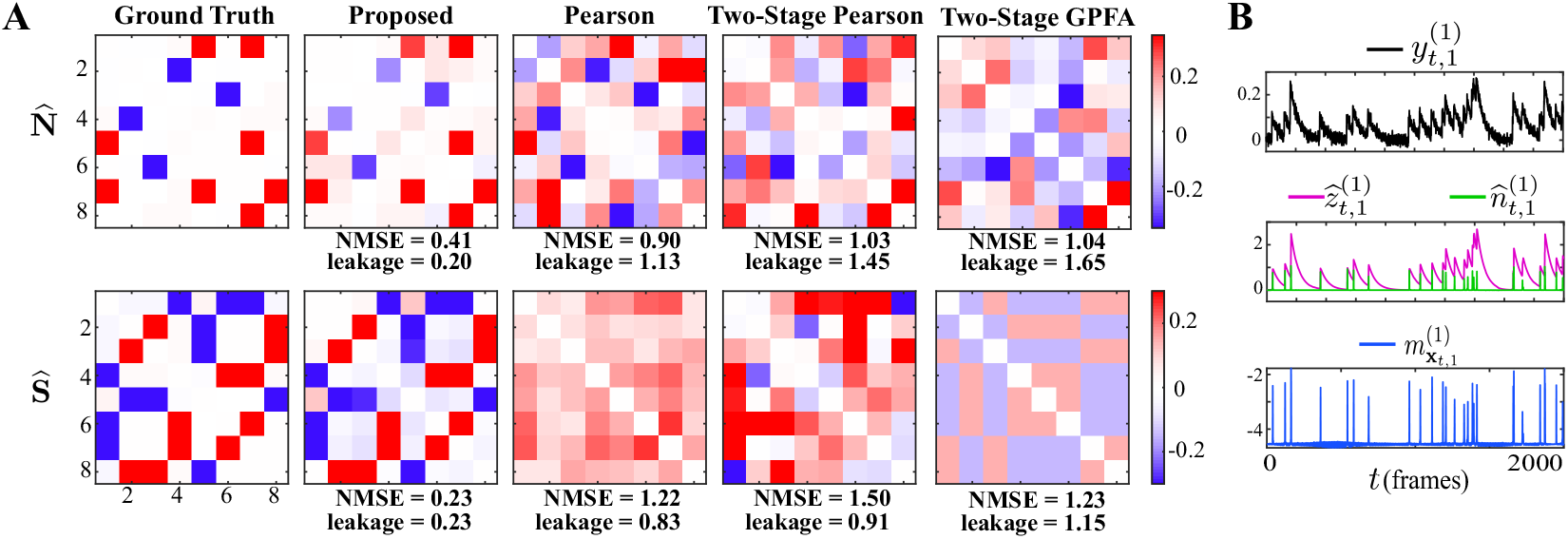
Results of simulation study 1. A) Estimated noise and signal correlation matrices from different methods. Rows from left to right: ground truth, proposed method, Pearson correlations from two-photon recordings, two-stage Pearson estimates and two-stage GPFA estimates. The normalized mean squared error (NMSE) of each estimate with respect to the ground truth and the leakage effect quantified by the ratio between out-of-network and in-network power (leakage) are indicated below each panel. B) Simulated fluorescence observations (black), estimated calcium concentrations (purple), putative spikes (green) and estimated mean of the latent state (blue) by the proposed method, for the first trial of neuron 1. **Figure 2-Figure supplement 1.** Sensitivity of two-stage estimates to the choice of the underlying spike deconvolution technique. **Figure 2-Figure supplement 2.** Performance of two-stage estimates based on ground truth spikes. **Figure 2-Figure supplement 3.** Proposed estimates based on simulated data with model mismatch and at lower SNR.

The first column of ***Figure 2***-A shows the ground truth noise (top) and signal (bottom) correlations (diagonal elements are all equal to 1 and omitted for visual convenience). The second column shows estimates of the noise and signal correlations using our proposed method, which closely match the ground truth. The third, fourth and fifth columns, respectively, show the results of the Pearson correlations from the two-photon data, two-stage Pearson, and two-stage GPFA estimation methods. Through a qualitative visual inspection, it is evident that these methods incur high false alarms and mis-detections of the ground truth correlations.

To quantify these comparisons, the normalized mean square error (NMSE) of different estimates with respect to the ground truth are shown below each of the subplots (***Figure 2***-A). Our proposed method achieves the lowest NMSE compared to the others. Furthermore, we observed a significant mixing between signal and noise correlations in these other estimates. To quantify this leakage effect, we first classified each of the correlation entries as in-network or out-of-network, based on being non-zero or zero in the ground truth, respectively (see Methods and Materials). We then computed the ratio between the power of out-of-network components and the power of in-network components as a measure of leakage. The leakage ratios are also reported in ***Figure 2***-A. The leakage of our proposed estimates is the lowest of all four techniques, in estimating both the signal and noise correlations. In order to further probe the performance of our proposed method, the simulated observations 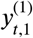, estimated calcium concentration 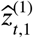, the putative spikes 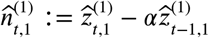, and the estimated mean of the latent state 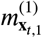, for the first trial of the first neuron are shown in ***Figure 2***-B. These results demonstrate the ability of the proposed estimation framework in accurately identifying the latent processes, which in turn leads to an accurate estimation of the signal and noise correlations as shown in ***Figure 2***-B.

The main sources of the observed performance gap between our proposed method and the existing ones are the bias incurred by treating the fluorescence traces as spikes, low spiking rates, non-linearity of spike generation with respect to intrinsic and external covariates, and sensitivity to spike deconvolution. For the latter, we demonstrated the sensitivity of the two-stage Pearson estimates to the choice of the deconvolution technique in ***Figure 2-Figure Supplement 1***. Furthermore, in order to isolate the effect of said non-linearities on the estimation performance, we applied the two-stage methods to ground truth spikes in ***Figure 2-Figure Supplement 2***. Our analysis showed that both two-stage estimates incur significant estimation errors even if the spikes were recovered perfectly, mainly due to the limited number of trials (*L* = 20 here). In accordance with our theoretical analysis of the asymptotic behavior of the conventional signal and noise correlation estimates given in ***Appendix 1***, we also showed in ***Figure 2-Figure Supplement 2*** that the performance of the two-stage Pearson estimates based on ground truth spikes, but using *L* = 1000 trials, dramatically improves. Our proposed method, however, was capable of producing reliable estimates with the number of trials as low as *L* = 20, which is typical in two-photon imaging experiments.

Finally, since real data does not necessarily follow the proposed generative model, to test the robustness of the proposed algorithm and modeling framework (with first-order autoregressive calcium dynamics assumption as outlined in Methods and Materials), we applied our method on simulated data generated based on a mismatched model (second-order autoregressive calcium dynamics), and at a lower signal-to-noise ratio (SNR) compared to the setting of ***Figure 2***. ***Figure 2-Figure Supplement 3*** shows the corresponding noise and signal correlations estimated by the proposed method under these conditions. Even though the performance slightly degrades (in terms of NMSE and leakage), our method is able to recover the underlying correlations faithfully under model mismatch and low SNR.

### Simulation study 2: Spontaneous activity

Next, we present the results of a simulation study in the absence of external stimuli (i.e. **s**_*t*_ = **0**), pertaining to the spontaneous activity condition. It is noteworthy that the proposed method can readily be applied to estimate noise correlations during spontaneous activity, by simply setting the external stimulus **s**_*t*_, and the receptive field **d**_*j*_ to zero in the update rules (see Methods and Materials for details). We simulated the ensemble spiking activity based on a Poisson process (***Smith and Brown, 2003***) using a discrete time-rescaling procedure (***Brown et al., 2002***; ***Smith and Brown, 2003***), so that the data are generated using a different model than that used in our inference framework (i.e., Bernoulli process with a logistic link as outlined in Methods and Materials). As such, we eliminated potential performance biases in favor of our proposed method by introducing the aforementioned model mismatch. We simulated *L* = 20 independent trials of spontaneous activity of *N* = 30 neurons, observed for a time duration of *T* = 5000 time frames. The number of neurons in this study is notably larger than that used in the previous one, to examine the scalability of our proposed approach with respect to the ensemble size.

***Figure 3*** shows the comparison of the noise correlation matrices estimated by our proposed method, Pearson correlations from two-photon recordings, two-stage Pearson, and two-stage GPFA estimates, with respect to the ground truth. The Pearson and the two-stage estimates are highly variable and result in excessive false detections. Our proposed estimate, however, closely follows the ground truth, which is also reflected by the comparatively lower NMSE and leakage ratios, in spite of the mismatch between the models used for data generation and inference. It is noteworthy that the proposed method exhibits favorable scaling with respect to the ensemble size, thanks to the underlying low-complexity variational updates (see Methods and Materials).

**Figure 3.**
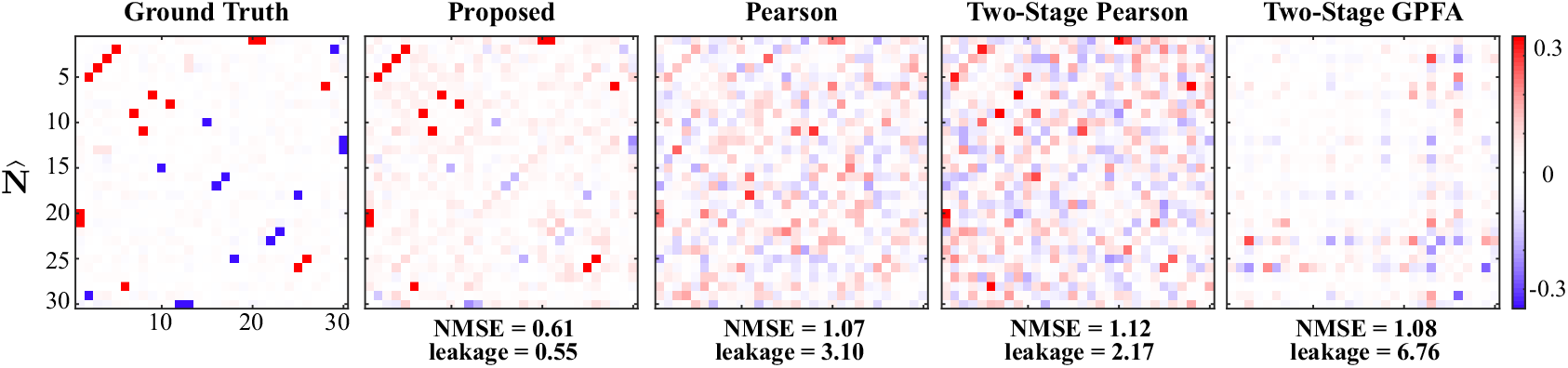
Results of simulation study 2. Estimated noise correlation matrices using different methods based from spontaneous activity data. Rows from left to right: ground truth, proposed method, Pearson correlations from two-photon recordings, two-stage Pearson and two-stage GPFA estimates. The normalized mean squared error (NMSE) of each estimate with respect to the ground truth and the ratio between out-of-network power and in-network power (leakage) are shown below each panel.

### Real data study 1: Mouse auditory cortex under random tone presentation

We next applied our proposed method to experimentally recorded two-photon observations from the mouse primary auditory cortex (A1). The dataset consisted of recordings from 371 excitatory neurons in layer 2/3 A1, from which we selected *J* = 16 neurons which exhibited the highest level of activity. A random sequence of four tones was presented to the mouse, with the same sequence being repeated for *L* = 10 trials. Each trial consisted of *T* = 3600 time frames, and each tone was two seconds long followed by a four-second silent period (see Methods and Materials for details). The comparison of the noise and signal correlation estimates obtained by our proposed method, Pearson correlations from two-photon recordings, two-stage Pearson and two-stage GPFA methods is shown in ***Figure 4***-A. The spatial map of the 16 neurons considered in the analysis in the field of view is shown in ***Figure 4***-B. ***Figure 4***-C shows the stimulus tone sequence *s_t_*, two-photon observations 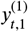, estimated calcium concentration 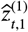, the putative spikes 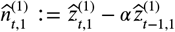 and the estimated mean of the latent state 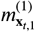, for the first trial of the first neuron.

**Figure 4.**
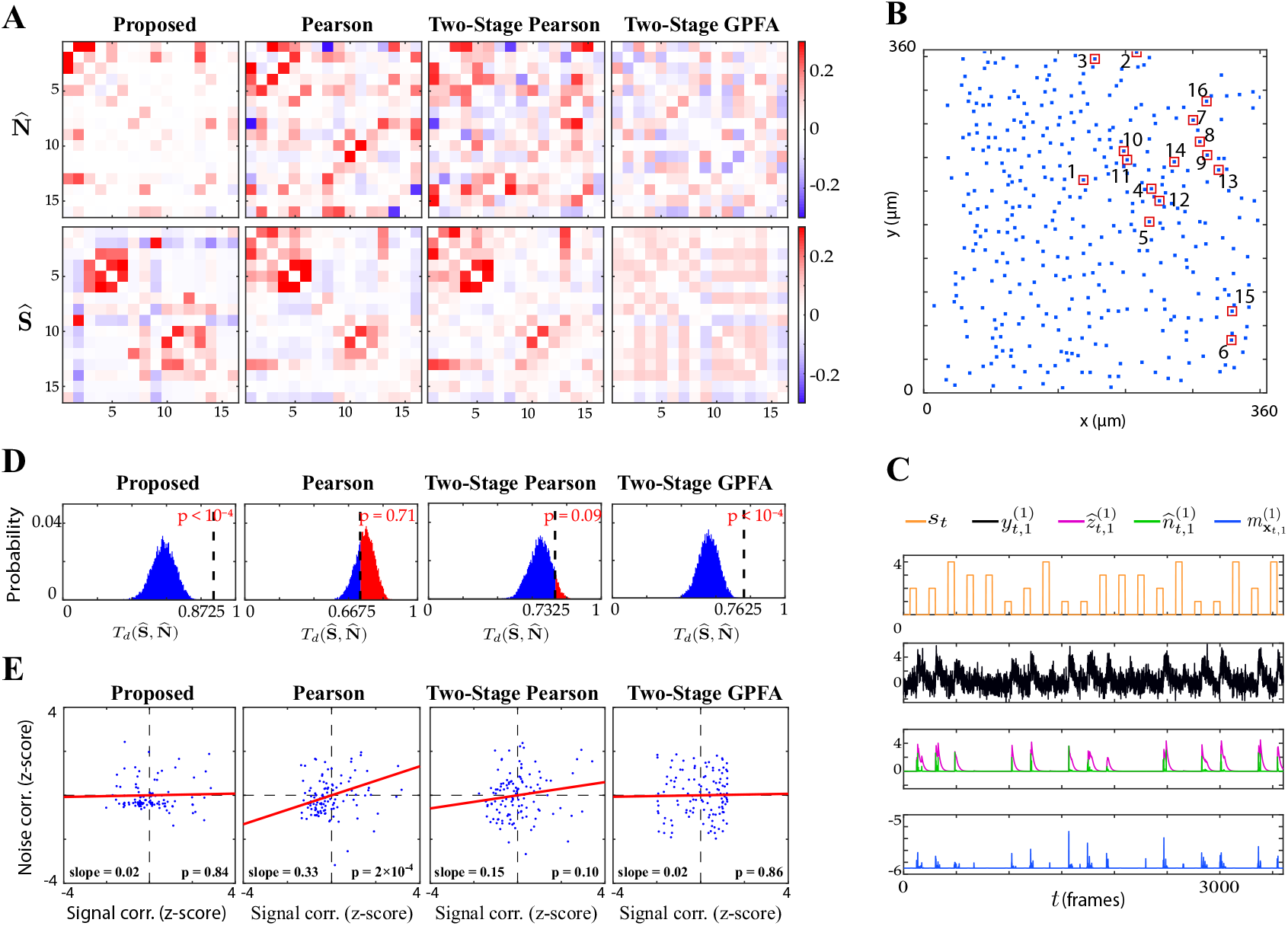
Application to experimentally-recorded data from the mouse A1. A) Estimated noise (top) and signal (bottom) correlation matrices using different methods. Rows from left to right: proposed method, Pearson correlations from two-photon data, two-stage Pearson and two-stage GPFA estimates. B) Location of the selected neurons with the highest activity in the field of view. C) Presented tone sequence (orange), observations (black), estimated calcium concentrations (purple), putative spikes (green) and estimated mean latent state (blue) in the first trial of the first neuron. D) Null distributions of chance occurrence of dissimilarities between signal and noise correlation estimates using different methods. The observed test statistic in each case is indicated by a dashed vertical line. E) Scatter plots of signal vs. noise correlations for individual cell pairs (blue dots) corresponding to each method. Data were normalized for comparison by computing z-scores. For each case, the linear regression model fit is shown in red, and the slope and p-value of the t-test are indicated as insets.

We estimated the Best Frequency (BF) of each neuron as the tone that resulted in the highest level of fluorescence activity. The results in ***Figure 4***-A are organized such that the neurons with the same BF are neighboring, with the BF increasing along the diagonal. Thus, expectedly (***Bowen et al., 2020***) our proposed method as well as the Pearson and two-stage Pearson estimates show high signal correlations along the diagonal. However, the two-stage GPFA estimates do not reveal such a structure. By visual inspection, as also observed in the simulation studies, the Pearson correlations from two-photon recordings, two-stage Pearson and two-stage GPFA estimates have significant leakage between the signal and noise correlations, whereas our proposed signal and noise correlation estimates in ***Figure 4***-A suggest distinct spatial structures.

To quantify this visual comparison, we used a statistic based on the Tanimoto similarity metric (***Lipkus, 1999***), denoted by *T_s_*(**X, Y**) for two matrices **X** and **Y**. As a measure of dissimilarity, we used *T_d_*(**X, Y**) := 1 – *T_s_*(**X, Y**) (see Methods and Materials). The comparison of 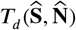 for the four estimates is presented in the second column of ***Table 1***. To assess statistical significance, for each comparison we obtained null distributions corresponding to chance occurrence of dissimilarities using a shuffling procedure as shown in ***Figure 4***-D, and then computed one-tailed *p*-values from those distributions (see Methods and Materials for details). ***Table 1*** and ***Figure 4***-D includes these *p*-values, which show that the proposed estimates (boldface numbers in ***Table 1***, second column) indeed have the highest dissimilarity between signal and noise correlations. The higher leakage effect in the other three estimates is also reflected in their smaller 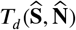 values. To further investigate this effect, we have depicted the scatter plots of signal vs. noise correlations estimated by each method in ***Figure 4***-E. To examine the possibility of the leakage effect on a pairwise basis, we performed linear regression in each case. The slope of the model fit, the p-value for the corresponding t-test, and the R^2^ values are reported in the third and fourth columns of ***Table 1*** (the slope and p-values are also shown as insets in ***Figure 4***-E). Consistent with the results of ***Winkowski and Kanold*** (***2013***), the Pearson estimates suggest a significant correlation between the signal and noise correlation pairs (as indicated by the higher slope in ***Figure 4***-E). However, it is noteworthy that none of the other estimates (including the proposed estimates) in ***Figure 4***-E register a significant trend between signal and noise correlations. This further corroborates our assessment of the high leakage between signal and noise correlations in Pearson estimates, since such a leakage effect could result in overestimation of the trend between the signal and noise correlation pairs. It is noteworthy that the signal and noise correlations estimated by our proposed method show no pairwise trend, suggesting distinct patterns of stimulus-dependent and stimulus-independent functional connectivity.

**Table 1.** Dissimilarity metric statistics for the estimates in ***Figure 4***-A (also illustrated in ***Figure 4***-D), linear regression statistics of the comparison between signal and noise correlations in ***Figure 4***-E, and the average NMSE across 50 trials used in the shuffling procedure illustrated in ***Figure 5***-A.

| Estimate | Dissimilarity $T_d(\hat{S}, \hat{N})$<br>(**Figure 4-D**) | Regression statistics (**Figure 4-E**) | | Shuffling test (**Figure 5**) | |
| --- | --- | --- | --- | --- | --- |
| | | slope (p-value) | R<sup>2</sup> value | NMSE in $\hat{N}$ | NMSE in $\hat{S}$ |
| Proposed | **0.8725** ($p < 10^{-4}$) | 0.02 ($p = 0.84$) | $4 \times 10^{-4}$ | **$1.07 \pm 0.16$** | **$1.32 \pm 0.19$** |
| Pearson | 0.6675 ($p = 0.71$) | 0.33 ($p = 2 \times 10^{-4}$) | 0.11 | 0 | 0 |
| Two-Stage Pearson | 0.7325 ($p = 0.09$) | 0.15 ($p = 0.10$) | 0.02 | $1.84 \pm 0.34$ | $0.55 \pm 0.12$ |
| Two-Stage GPFA | 0.7625 ($p < 10^{-4}$) | 0.02 ($p = 0.86$) | $3 \times 10^{-4}$ | $2.32 \pm 0.52$ | $2.26 \pm 0.51$ |

Given that the ground truth correlations are not available for a direct comparison, we instead performed a test of specificity that reveals another key limitation of existing methods. Fluorescence observations exhibit structured dynamics due to the exponential intracellular calcium concentration decay (as shown in ***Figure 4***-C, for example), which are in turn related to the underlying spikes that are driven non-linearly by intrinsic/extrinsic stimuli as well as the properties of the indicator used. As such, an accurate inference method is expected to be specific to this temporal structure. To test this, we randomly shuffled the *T* time frames consistently in the same order in all trials, in order to fully break the temporal structure governing calcium decay dynamics, and then estimated correlations from these shuffled data using the different methods. The resulting estimates of noise correlations are shown in ***Figure 5***-A for one instance of such shuffled data. The average NMSE for a total of 50 shuffled samples with respect to the original un-shuffled estimates (in ***Figure 4***-A) are tabulated in the fifth and sixth columns of ***Table 1***, and are also indicated below each panel in ***Figure 5***-A.

**Figure 5.**
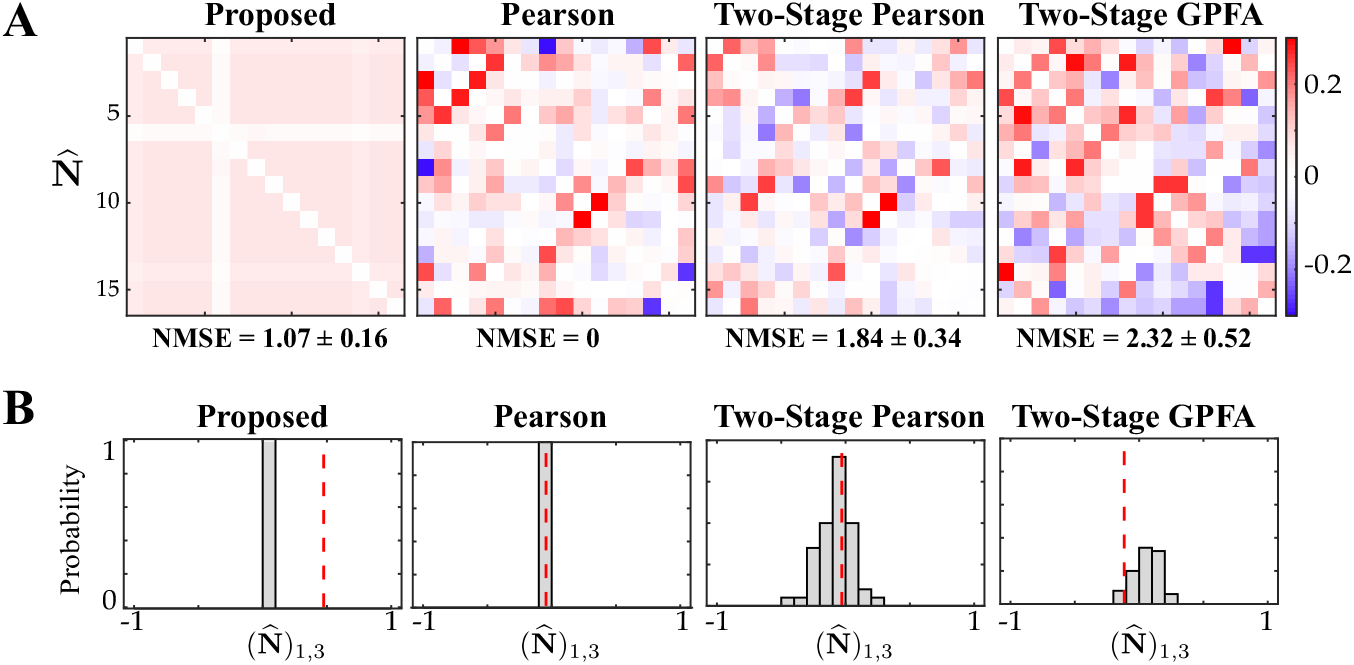
Assessing the specificity of different estimation results shown in ***Figure 4***. Rows from left to right: proposed method, Pearson correlations from two-photon data, two-stage Pearson and two-stage GPFA estimates. A) The estimated noise correlations using different methods after random temporal shuffling of the observations. The mean and standard deviation of the NMSE across 50 trials are indicated below each panel. B) Histograms of the noise correlation estimates between the first and third neurons over the 50 temporal shuffling trials. The estimate based on the original (un-shuffled) data in each case is indicated by a dashed vertical line.

A visual inspection of ***Figure 5***-A shows that the Pearson correlations from two-photon recordings expectedly remain unchanged. Since this method treats each time frame to be independent, temporal shuffling does not impact the correlations in anyway. On the other extreme, both of the two-stage estimates seem to detect highly variable and large correlation values, despite operating on data that lacks any relevant temporal structure. Our proposed method, however, remarkably produces negligible correlation estimates. Although both the two-stage and proposed estimates show variability with respect to the shuffled data (***Table 1***), the standard deviation of the NMSE values of our proposed method are considerably smaller than those of the two-stage methods (***Table 1***, fifth column). For further inspection, the histograms of a single element 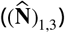 of the estimated correlation matrices across the 50 shuffling trials are shown in ***Figure 5***-B. The original un-shuffled estimates are marked by the dashed vertical lines in each case. The proposed estimate in ***Figure 5***-B is highly concentrated around zero, even though the un-shuffled estimate is non-zero. However, the two-stage estimates produce correlations that are widely variable across the shuffling trials. This analysis demonstrates that our proposed method is highly specific to the temporal structure of fluorescence observations, whereas the Pearson correlations from two-photon recordings, two-stage Pearson and two-stage GPFA methods fail to be specific.

### Real data study 2: Spontaneous vs. stimulus-driven activity in the mouse A1

To further validate the utility of our proposed methodology, we applied it to another experimentally-recorded dataset from the mouse layer 2/3 A1. This experiment pertained to trials of presenting a sequence of short white noise stimuli, randomly interleaved with silent trials of the same duration. The two-photon recordings thus contained episodes of stimulus-driven and spontaneous activity (see Methods and Materials for details). Under these experimental conditions, it is expected that the noise correlations are invariant across the spontaneous and stimulus-driven conditions, and that the signal and noise correlation patterns are distinct (***Kohn et al., 2016***; ***Montijn et al., 2014***; ***Rothschild et al., 2010***; ***Keeley et al., 2020***). Each trial consisted of *T* = 765 frames. We selected *N* = 10 neurons with the highest level of activity for the analysis, each with *L* = 10 trials.

***Figure 6***-A shows the resulting noise and signal correlation estimates under the spontaneous (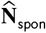, top) and stimulus-driven (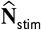 and 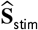, bottom) conditions. ***Figure 6***-B shows the spatial map of the 10 neurons considered in the analysis in the field of view. A visual inspection of the first column of ***Figure 6***-A indeed suggests that 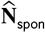 and 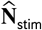 are saliently similar, and distinct from 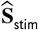. The Pearson correlations obtained from two-photon data (second column) and the two-stage Pearson and GPFA estimates (third and fourth columns, respectively), however, evidently lack this structure. As in the previous study, we quantified this visual comparison using the similarity metric *T_s_*(**X, Y**) and the dissimilarity metric *T_d_*(**X, Y**) (see Methods and Materials for details). These statistics are reported in ***Table 2*** along with the *p*-values (null distributions are shown in ***Figure 6-Figure Supplement 1***), which show that the only significant outcomes (boldface numbers) are those of our proposed method.

**Figure 6.**
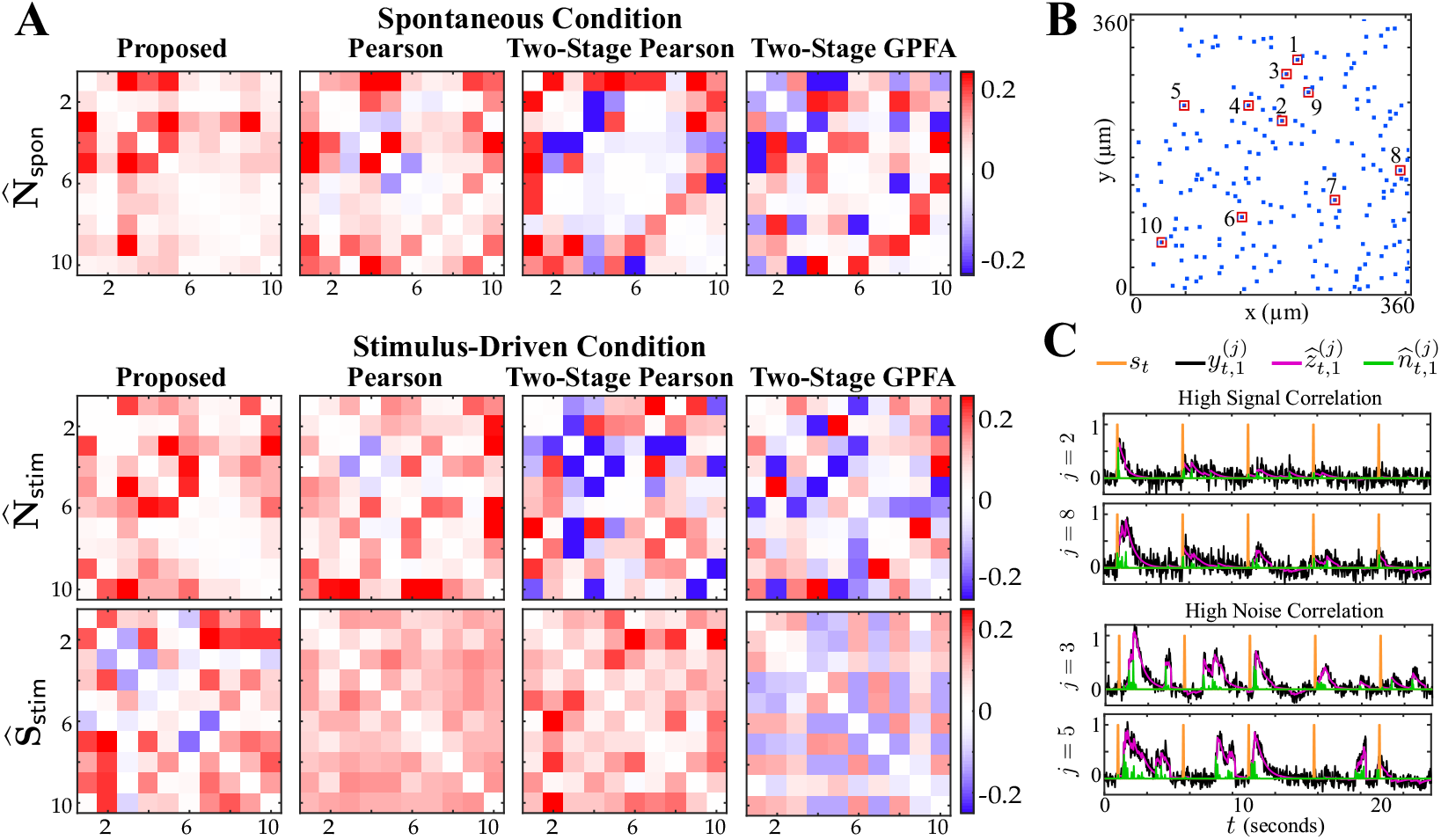
Comparison of spontaneous and stimulus-driven activity in the mouse A1. A) Estimated noise and signal correlation matrices under spontaneous (top) and stimulus-driven (bottom) conditions. Rows from left to right: proposed method, Pearson correlations from two-photon data, two-stage Pearson and two-stage GPFA estimates. B) Location of the selected neurons with highest activity in the field of view. C) Stimulus onsets (orange), observations (black), estimated calcium concentrations (purple) and putative spikes (green) for the first trial from two pairs of neurons with high signal correlation (top) and high noise correlation (bottom), as identified by the proposed estimates. **Figure 6-Figure supplement 1.** Histograms of the similarity/dissimilarity metrics under the shuffling procedure.

**Table 2.** Similarity/dissimilarity metric statistics for the estimates in ***Figure 6***.

| Estimation Method | $T_s(\hat{\mathbf{N}}_{\text{spon}}, \hat{\mathbf{N}}_{\text{stim}})$ | $T_d(\hat{\mathbf{S}}_{\text{stim}}, \hat{\mathbf{N}}_{\text{stim}})$ |
| --- | --- | --- |
| Proposed | **0.5518** ($p = 0.01$) | **0.7625** ($p = 0.01$) |
| Pearson | 0.3031 ($p = 0.61$) | 0.5025 ($p = 0.92$) |
| Two-Stage Pearson | 0.2790 ($p = 0.05$) | 0.7925 ($p = 0.39$) |
| Two-Stage GPFA | 0.2008 ($p = 0.50$) | 0.7825 ($p = 0.22$) |

Furthermore, ***Figure 6***-C shows the time course of the stimulus, observations, estimated calcium concentrations and putative spikes for the first trial from two pairs of neurons with high signal correlation (*j* = 2,8, top) and high noise correlation (*j* = 3,5, bottom). As expected, the putative spiking activity of the neurons with high signal correlation (top) are closely time-locked to the stimulus onsets. The activity of the two neurons with high noise correlation (bottom), however, is not time-locked to the stimulus onsets, even though the two neurons exhibit highly correlated activity. The correlations estimated via the proposed method thus encode substantial information about the inter-dependencies of the spiking activity of the neuronal ensemble.

### Real data study 3: Spatial analysis of signal and noise correlations in the mouse A1

Lastly, we applied our proposed method to examine the spatial distribution of signal and noise correlations in the mouse A1 layers 2/3 and 4 (data from ***Bowen et al.*** (***2020***)). The dataset included fluorescence activity recorded during multiple experiments of presenting sinusoidal amplitude-modulated tones, with each stimulus being repeated across several trials (see Methods and Materials and ***Bowen et al.*** (***2020***) for experimental details). In each experiment, we selected around 20 neurons with highest spiking rates for the subsequent analysis. For brevity, we compare the estimates of signal and noise correlations using our proposed method only with those obtained by Pearson correlations from the two-photon data. The latter method was also used in previous analyses of data from this experimental paradigm (***Winkowski and Kanold, 2013***).

In parallel to the results reported in ***Winkowski and Kanold*** (***2013***), ***Figure 7***-A and ***Figure 7***-B illustrate the correlation between the signal and noise correlations in layers 2/3 and 4, respectively. Consistent with the results of ***Winkowski and Kanold*** (***2013***), the signal and noise correlations exhibit positive correlation in both layers, regardless of the method used. However, the correlation coefficients (i.e., slopes in the insets) identified by our proposed method are notably smaller than those obtained from Pearson correlations, in both layer 2/3 (***Figure 7***-A) and layer 4 (***Figure 7***-B). Comparing this result with our simulation studies suggests that the stronger linear trend between the signal and noise correlations observed using the Pearson correlation estimates is likely due to the mixing between the estimates of signal and noise correlations. As such, our method suggests that the signal and noise correlations may not be as highly correlated with one another as indicated in previous studies of layer 2/3 and 4 in mouse A1.

**Figure 7.**
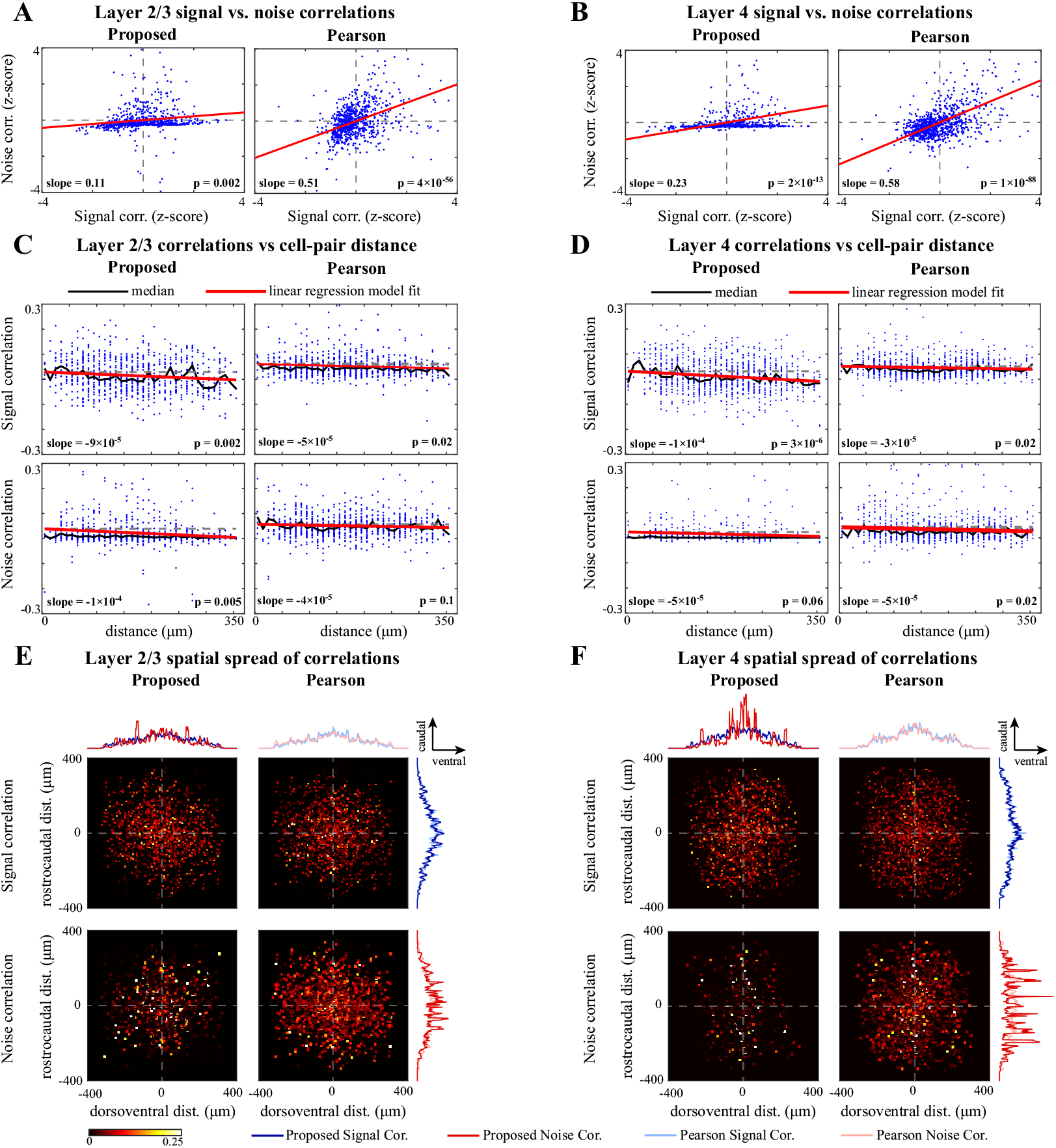
Comparison of signal and noise correlations across layers 2/3 and 4. A) Scatter-plot of noise vs. signal correlations (blue) for individual cell-pairs in layer 2/3, based on the proposed (left) and Pearson estimates (right). Data were normalized for comparison by computing z-scores. The linear model fits are shown in red, and the slope and p-value of the t-tests are indicated as insets. Panel B corresponds to layer 4 in the same organization as panel A. C) Signal (top) and noise (bottom) correlations vs. cell-pair distance in layer 2/3, based on the proposed (left) and Pearson estimates (right). Distances were binned to 10 *μm* intervals. The median of the distributions (black) and the linear model fit (red) are shown in each panel. The slope of the linear model fit, and the p-value of the t-test are also indicated as insets. Dashed horizontal lines indicate the zero-slope line for ease of visual comparison. Panel D corresponds to layer 4 in the same organization as panel C. E) Spatial spread of signal (top) and noise (bottom) correlations in layer 2/3, based on the proposed (left) and Pearson estimates (right). The horizontal and vertical axes in each panel respectively represent the relative dorsoventral and rostrocaudal distances between each cell-pair, and the heat-map indicates the magnitude of correlations. Marginal distributions of the signal (blue) and noise (red) correlations along the dorsoventral and rostrocaudal axes for the proposed method (darker colors) and Pearson method (lighter colors) are shown at the top and right side of the sub-panels. Panel F corresponds to layer 4 in the same organization as panel E. **Figure 7-Figure supplement 1.** Comparing the marginal distributions of signal and noise correlations along the dorsoventral and rostrocaudal axes. **Figure 7-Figure supplement 2.** Marginal angular distributions of signal and noise correlations.

Next, to evaluate the spatial distribution of signal and noise correlations, we plotted the correlation values for pairs of neurons as a function of their distance for layer 2/3 (***Figure 7***-C) and layer 4 (***Figure 7***-D). The distances were discretized using bins of length 10 *μm.* The scatter of the correlations along with their median at each bin are shown in all panels. Then, to examine the spatial trend of the correlations, we performed linear regression in each case. The slope of the model fit, the p-value for the corresponding t-test, and the R^2^ values are reported in ***Table 3*** (the slope and p-values are also shown as insets in ***Figure 7***-C & D).

**Table 3.** Linear regression statistics for the analysis of correlations vs. cell-pair distance

| Correlations | Statistics of layer 2/3 correlations |  | Statistics of layer 4 correlations |  |
| --- | --- | --- | --- | --- |
|  | slope (p-value) | R<sup>2</sup> value | slope (p-value) | R<sup>2</sup> value |
| Proposed Signal Corr. | **$-9 \times 10^{-5}$** ($p = 0.002$) | 0.012 | **$-1 \times 10^{-4}$** ($p = 3 \times 10^{-6}$) | 0.023 |
| Pearson Signal Corr. | $-5 \times 10^{-5}$ ($p = 0.02$) | 0.007 | $-3 \times 10^{-5}$ ($p = 0.02$) | 0.005 |
| Proposed Noise Corr. | **$-1 \times 10^{-4}$** ($p = 0.005$) | 0.010 | **$-5 \times 10^{-5}$** ($p = 0.06$) | 0.004 |
| Pearson Noise Corr. | $-4 \times 10^{-5}$ ($p = 0.1$) | 0.003 | $-5 \times 10^{-5}$ ($p = 0.02$) | 0.005 |

From ***Table 3*** and ***Figure 7***-C & D (upper panels), it is evident that the signal correlations show a significant negative trend with respect to distance, using both methods and in both layers. However, the slope of these negative trends identified by our method (boldface numbers in ***Table 3***) is notably steeper than those identified by Pearson correlations. On the other hand, the trends of the noise correlations with distance (bottom panels) are different between our proposed method and Pearson correlations: our proposed method shows a significant negative trend in layer 2/3, but not in layer 4, whereas the Pearson correlations of the two-photon data suggest a significant negative trend in layer 4, but not in layer 2/3. In addition, the slopes of these negative trends identified by our method (boldface numbers in ***Table 3***) are steeper than or equal to those identified by Pearson correlations.

Our proposed estimates indicate that noise correlations are sparser and less widespread in layer 4 (***Figure 7***-D) than in layer 2/3 (***Figure 7***-C). To further investigate this observation, we depicted the two-dimensional spatial spread of signal and noise correlations in both layers and for both methods in ***Figure 7***-E & F, by centering each neuron at the origin and overlaying the individual spatial spreads. The horizontal and vertical axes in each panel respectively represent the relative dorsoventral and rostrocaudal distances, and the heat-maps represent the magnitude of correlations. Comparing the proposed noise correlation spread in ***Figure 7***-E with the corresponding spread in ***Figure 7***-F, we observe that the noise correlations in layer 2/3 are indeed more widespread and abundant than in layer 4.

It is notable that the spatial spreads of signal and noise correlations based on the Pearson estimates are remarkably similar in both layers (***Figure 7***-E & F, right panels), whereas they are saliently different for our proposed estimates (***Figure 7***-E & F, left panels). This further corroborates our hypothesis on the possibility of high mixing between the signal and noise correlation estimates obtained by the Pearson correlation of two-photon data. To further examine the differences between the signal and noise correlations, the marginal distributions along the dorsoventral and rostrocaudal axes are shown in ***Figure 7***-E & F, selectively overlaid for ease of visual comparison. To quantify the differences between the spatial distributions of signal and noise correlations estimated by each method, we performed Kolmogorov-Smirnov (KS) tests on each pair of marginal distributions, which are summarized in ***Figure 7-Figure Supplement 1***. Although the marginal distributions of signal and noise correlations are significantly different in all cases from both methods, the effect sizes of their difference (KS statistics) are notably higher for our proposed estimates compared to those of the Pearson estimates.

Finally, it is noteworthy that the spatial spreads of correlations for either method and in each layer suggest non-uniform angular distributions with possibly directional bias. To test this effect, we computed the angular marginal distributions and performed KS tests for non-uniformity, which are reported in ***Figure 7-Figure Supplement 2***. These tests indicate that all distributions are significantly non-uniform. In addition, the angular distributions of both signal and noise correlations in layer 4 exhibit salient modes in the rostrocaudal direction, whereas they are less directionally selective in layer 2/3 (***Figure 7-Figure Supplement 2***).

In summary, the spatial trends identified by our proposed method are consistent with empirical observations of spatially heterogeneous pure-tone frequency tuning by individual neurons in auditory cortex (***Winkowski and Kanold, 2013***). The improved correspondence of our proposed method compared to results obtained using Pearson correlations could be the result of the demixing of signal and noise correlations in our method. As a result of the demixing, our proposed method also suggests that noise correlations have a negative trend with distance in layer 2/3, but are much sparser and spatially flat in layer 4. In addition, the spatial spread patterns of signal and noise correlations are more structured and remarkably more distinct for our proposed method than those obtained by the Pearson estimates.

### Theoretical analysis of the bias and variance of the proposed estimators

Finally, we present a theoretical analysis of the bias and variance of the proposed estimator. Note that our proposed estimation method has been developed as a scalable alternative to the intractable maximum likelihood (ML) estimation of the signal and noise covariances (see Methods and Materials). In order to benchmark our estimates, we thus need to evaluate the quality of said ML estimates. To this end, we derived bounds on the bias and variance of the ML estimators of the kernel **d**_*j*_ for *j* = 1,∞, *N* and the noise covariance **∑**_*x*_. In order to simplify the treatment, we posit the following mild assumptions:

#### Assumption (1).

We assume a scalar time-varying external stimulus (i.e. **s**_*t*_ = *s_t_*, and hence **d**_*j*_ = *d_j_,* **d** = [*d*_1_*,d*_2_, ∞, *d_N_*]^τ^). Furthermore, we set the observation noise covariance to be 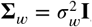, for notational convenience.

#### Assumption (2).

We derive the performance bounds in the regime where *T* and *L* are large, and thus do not impose any prior distribution on the correlations, which are otherwise needed to mitigate overfitting (see Methods and Materials).

#### Assumption (3).

We assume the latent noise process and stimulus to be slowly varying signals, and thus adopt a piece-wise constant model in which these processes are constant within consecutive windows of length *W* (i.e., **x**_*t,l*_ = ***x***_*W*_*k*_,*l*_ and *s_t_* = *s_W_k__*, for (*k* – 1)*W* + 1 ≤ *t* < *kW* and *k* = 1,∞,*K* with *W_k_* = (*k* – 1)*W* + 1 and *KW* = *T*) for our theoretical analysis, as is usually done in spike count calculations for conventional noise correlation estimates.

Our main theoretical result is as follows:

#### Theorem 1

(Performance Bounds). *Let* 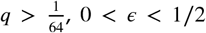, 0 < *ϵ* < 1/2, *and* 0 < *η* ≤ 1/2 *be fixed constants*, 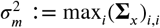 *and* 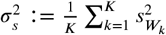. *Then, under Assumptions (1) - (3), the bias and variance of the maximum likelihood estimators* 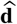 *and* 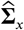, *conditioned on an event* 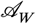 *with* 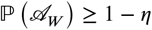 *satisfy*:

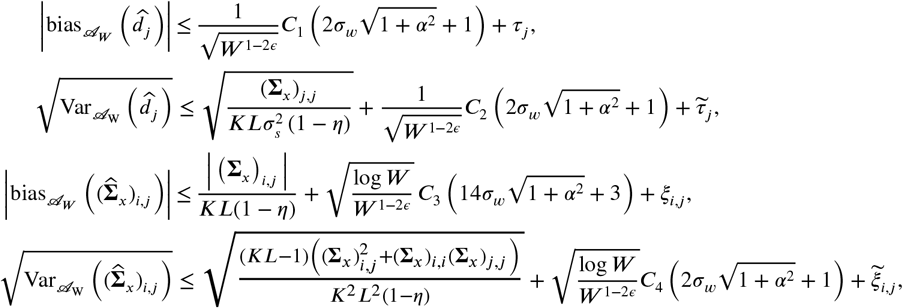

*for all i, j* = 1,2,∞, *N, if* 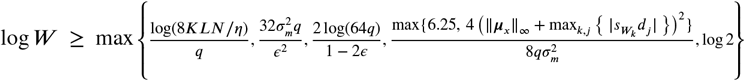, where *τ_j_* and 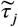 *denote bounded terms that are* 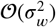 *or* 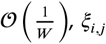 and 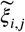 *denote bounded terms that are* 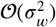 *or* 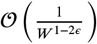 *and C*_1_, *C*_2_, *C*_3_ *and C*_4_ *are bounded constants given in* **Appendix 2**.

*Proof.* The proof of Theorem ***1*** is provided in ***Appendix 2.***

In order to discuss the implications of this theoretical result, several remarks are in order:

#### Remark 1: Achieving near oracle performance

A common benchmark in estimation theory is the performance of the idealistic *oracle* estimator, in which an oracle directly observes the true latent process ***x***_*t,l*_ and the true kernel *d_j_* and forms the correlation estimates. In this case, the oracle would incur zero bias and variance of order 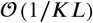 in estimating *d_j_*, and outputs an estimate of **∑**_*x*_ with bias and variance in the order of 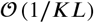. Theorem 1 indeed states that for sufficiently large *W* and small *σ_w_,* the bias and variance of the ML estimators are arbitrarily close to those of the oracle estimator. Recall that our variational inference framework is in fact a solution technique for the regularized ML problem. Hence, the bounds in Theorem 1 provide a benchmark for the expected performance of the proposed estimators, by quantifying the excess bias and variance over the performance of the oracle estimator.

#### Remark 2: Effect of the observation noise and observation duration

As the assumed window of stationarity *W* → ∞ (and hence the observation duration *T* → ∞), the loss of performance of the proposed estimators only depends on 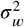, the variance of the observation noise. As a result, at a given observation noise variace 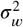, these bounds provide a sufficient upper bound on the time duration of the observations required for attaining a desired level of estimation accuracy. It is noteworthy that 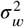 is typically small in practice, as it pertains to the effective observation noise and is significantly diminished by pixel averaging of the fluorescence traces following cell segmentation.

#### Remark 3: Effect of the number of trials

Finally, note that the bounds in Theorem 1 have terms that also drop as the number of trials *L* grows. These terms in fact pertain to the performance of the oracle estimator. As the number of trials grows (*L* → ∞), the oracle estimates become arbitrarily close to the true parameters **∑**_*x*_ and **d**_*j*_. Thus, our theoretical performance bounds also provide a sufficient upper bound on the number of trials *L* required for the oracle estimator to attain a desired level of estimation accuracy.

## Discussion

We developed a novel approach for the joint estimation of signal and noise correlations of neuronal activities directly from two-photon calcium imaging observations and tested our method with experimental data. Existing widely used methods either take the fluorescence traces as surrogates of spiking activity, or first recover the unobserved spikes using deconvolution techniques, both followed by computing Pearson correlations or connectivity matrices. As such, they typically result in estimates that are highly biased and are heavily dependent on the choice of the spike deconvolution technique. We addressed these issues by using data with multiple repeated trials and explicitly relating the signal and noise covariances to the observed two-photon data via a multi-tier Bayesian model that accounts for the observation process and non-linearities involved in spiking activity. We developed an efficient estimation framework by integrating techniques from variational inference and state-space estimation. We established performance bounds on the bias and variance of the proposed estimators, which revealed favorable scaling with respect to the observation noise and trial length.

We demonstrated the utility of our proposed estimation framework on both simulated and experimentally-recorded data from the mouse auditory cortex. Our analysis showed that, unlike the aforementioned methods, our estimates provide noise correlation structures that are highly invariant across spontaneous and stimulus-driven conditions, while producing signal correlation structures that are largely distinct from those given by the noise correlation. These results provide evidence for the involvement of distinct functional neuronal network structures in encoding the stimulus-dependent and stimulus-independent information.

Our analysis of the relationship between the signal and noise correlations in layers 2/3 and 4 in mouse A1 indicates a smaller correlation between signal and noise correlations than previously reported (***Winkowski and Kanold, 2013***). Thus, our proposed method suggests that the signal and noise correlations reflect distinct circuit mechanisms of sound processing in layers 2/3 vs 4. The spatial distribution of signal correlations obtained by our method was consistent with previous work showing significant negative trends with distance (***Winkowski and Kanold, 2013***). However, in addition, our proposed method revealed a significant negative trend of noise correlations with distance in layer 2/3, but not in layer 4, in contrast to the outcome of Pearson correlation analysis. The lack of a negative trend in layer 4 could be attributed to the sparse nature of the noise correlation spread in layer 4, as revealed by our analysis of two-dimensional spatial spreads. The latter analysis indeed revealed that the noise correlations in layer 2/3 are more widespread than those in layer 4, consistent with existing work based on whole-cell patch recordings (***Meng et al., 2017a,b***). In addition, the two-dimensional spatial spreads of signal and noise correlations obtained by our method are more distinct than those obtained by Pearson correlations. The spatial spreads also allude to directionality of the functional connectivity patterns, with a notable rostrocaudal preference in layer 4.

It is noteworthy that the proposed method can scale up favorably to larger populations of neurons, thanks to the underlying low-complexity variational updates in the inference procedure. Due to its minimal dependence on training data, our estimation framework is also applicable to single-session analysis of two-photon data with limited number of trials and duration. Another useful byproduct of the proposed framework is gaining access to approximate posterior densities in closed-form, which allows further statistical analyses such as construction of confidence intervals. Our proposed methodology can thus be used as a robust and scalable alternative to existing approaches for extracting neuronal correlations from two-photon calcium imaging data.

A potential limitation of our proposed model is the assumption that there is at most one spiking event per time frame for each neuron, in light of the fact that typical two-photon imaging frame durations are in the range of 30-100 ms. Average spike rates of excitatory neurons in mouse A1 layers 2/3 and 4 are of the order of < 10 Hz (***Forli et al., 2018***) and thus our assumption is reasonable for the current study, although it might not be optimal during bursting activity. Even though this assumption can be made more precise by adopting a Poisson model, that would render closed-form variational density updates intractable. Furthermore, in the regime of extremely low spiking rate and high observation noise, the proposed forward model may fail to capture the underlying correlations faithfully, and the performance is expected to degrade to those of existing methods based on Pearson correlations. Nevertheless, our method addresses key limitations of conventional signal and noise correlation estimators that persist in high spiking rate and high SNR conditions.

Our proposed estimation framework can be used as groundwork for incorporating other notions of correlation such as the connected correlation function (***Martin et al., 2020***), and to account for non-Gaussian and higher-order structures arising from spatiotemporal interactions (***Kadirvelu et al., 2017***; ***Yu et al., 2011***). Other possible extensions of this work include leveraging variational inference beyond the mean-field regime ***Wang and Blei*** (***2013***), extension to time-varying correlations that underlie rapid task-dependent dynamics, and extension to non-linear models such as those parameterized by neural networks (***Aitchison et al., 2017***). In the spirit of easing reproducibility, a MATLAB implementation of our proposed method as well as the data used in this work are made publicly available (***Rupasinghe, 2020***; ***Rupasinghe et al., 2021***).

## Methods and Materials

### Proposed forward model

Suppose we observe fluorescence traces of *N* neurons, for a total duration of *T* discrete-time frames, corresponding to *L* independent trials of repeated stimulus. Let 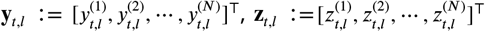, and 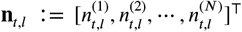 be the vectors of noisy observations, intracellular calcium concentrations, and ensemble spiking activities, respectively, at trial *l* and frame *t*. We capture the dynamics of ***y***_*t*,*l*_ by the following state-space model:

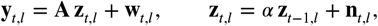

where **A** ∈ **ℝ**^*N×N*^ represents the scaling of the observations, **w**_*t,l*_ is zero-mean i.i.d. Gaussian noise with covariance **∑**_*w*_, and 0 ≤ *α* < 1 is the state transition parameter capturing the calcium dynamics through a first order model. Note that this state-space is non-Gaussian due to the binary nature of the spiking activity, i.e., 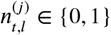. We model the spiking data as a point process or Generalized Linear Model with Bernoulli statistics (***Eden et al., 2004***; ***Paninski, 2004***; ***Smith and Brown, 2003***; ***Truccolo et al., 2005***):

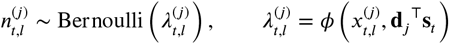

where 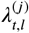 is the conditional intensity function (***Truccolo et al., 2005***), which we model as a non-linear function of the known external stimulus **s**_*t*_ and the other latent intrinsic and extrinsic trial-dependent covariates, 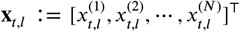. While we assume the stimulus **s**_*t*_ ∈ ℝ^*M*^ to be common to all neurons, we model the distinct effect of this stimulus on the *j*^th^ neuron via an unknown kernel **d**_*j*_ ∈ ℝ^*M*^, akin to the receptive field.

The non-linear mapping of our choice is the logistic link, which is also the canonical link for a Bernoulli process in the point process and Generalized Linear Model frameworks (***Truccolo et al., 2005***). Thus, we assume:

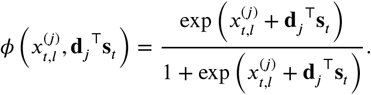

Finally, we assume the latent trial dependent covariates to be a Gaussian process 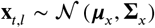, with mean 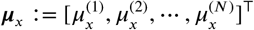 and covariance **∑**_*x*_.

The probabilistic graphical model in ***Figure 8*** summarizes the main components of the aforementioned forward model. According to this forward model, the underlying noise covariance matrix that captures trial-to-trial variability can be identified as **∑**_*x*_. The signal covariance matrix, representing the covariance of the neural activity arising from the repeated application of the stimulus **s**_*t*_, is given by **∑**_*s*_ := **D**^τ^ cov(**s**_*t*_. **s**_*t*_) **D**, where **D** := [**d**_1_, **d**_2_,∞, **d**_*N*_ ∈ ℝ^*M*×*M*^. The signal and noise correlation matrices, denoted by **S** and **N**, can then be obtained by standard normalization of **∑**_*s*_ and **∑**_*x*_:

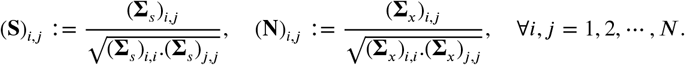

The main problem is thus to estimate {**∑**_*x*_. **D**} from the noisy and temporally blurred data 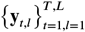.

**Figure 8.**
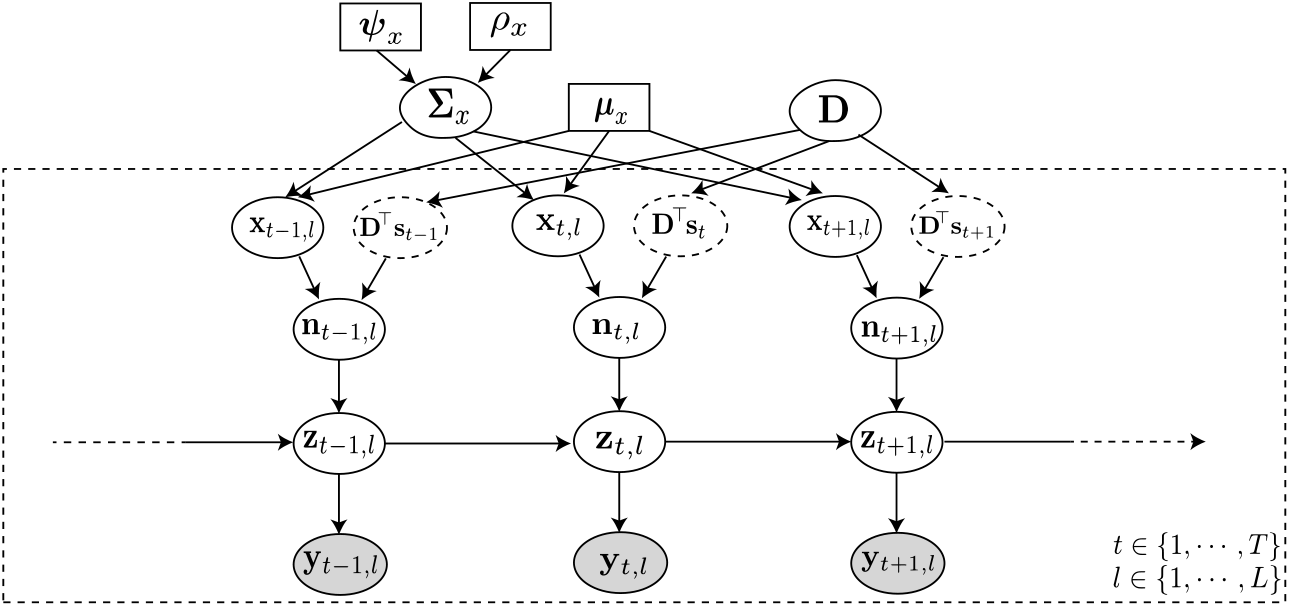
Probabilistic graphical model of the proposed forward model. The fluorescence observations at the *t*^th^ time frame and *l*^th^ trial: ***y**_t,l_*, are noisy surrogates of the intracellular calcium concentrations: **z**_*t,l*_. The calcium concentration at time *t* is a function of the spiking activity **n**_*t,l*_, and the calcium activity at the previous time point **z**_*t*–1,*l*_. The spiking activity is driven by two independent mechanisms: latent trial-dependent covariates **x**_*t,l*_, and contributions from the known external stimulus **s**_*t*_, which we model by **D**^τ^**s**_*t*_ (in which the receptive field **D** is unknown). Then, we model **x**_*t,l*_ as a Gaussian process with constant mean ***μ***_*x*_, and unknown covariance **∑**_*x*_. Finally, we assume the covariance **∑**_*x*_ to have an inverse Wishart prior distribution with hyper-parameters ***ψ***_*x*_ and *ρ_x_*. Based on this forward model, the inverse problem amounts to recovering the signal and noise correlations by directly estimating **∑**_*x*_ and **D** (top layer) from the fluorescence observations 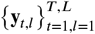 (bottom layer).

### Overview of the proposed estimation method

First, given a limited number of trials *L* from an ensemble with typically low spiking rates, we need to incorporate suitable prior assumptions to avoid overfitting. Thus, we impose a prior *p*_pr_(**∑**_*x*_) on the noise covariance, to compensate sparsity of data. A natural estimation method to estimate {**∑**_*x*_. **D**} in a Bayesian framework is to maximize the observed data likelihood 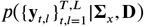, i.e., maximum likelihood (ML). Thus, we consider the joint likelihood of the observed data and latent processes to perform Maximum a Posteriori (MAP) estimation:

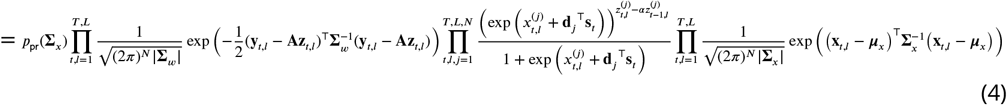

Inspecting this MAP problem soon reveals that estimating **∑**_*x*_ and **D** is a challenging task: 1) standard approaches such as Expectation-Maximization (EM) (***Shumway and Stoffer, 1982***) are intractable due to the complexity of the model, arising from the hierarchy of latent processes and the non-linearities involved in their mappings, and 2) the temporal coupling of the likelihood in the calcium concentrations makes any potential direct solver scale poorly with *T*.

Thus, we propose an alternative solution based on Variational Inference (VI) (***Beal, 2003***; ***Blei et al., 2017***; ***Jordan et al., 1999***). VI is a method widely used in Bayesian statistics to approximate unwieldy posterior densities using optimization techniques, as a low-complexity alternative strategy to Markov Chain Monte Carlo sampling (***Hastings, 1970***) or empirical Bayes techniques such as EM. To this end, we treat 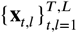 and **∑**_*x*_ as latent variables and 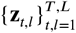 and **D** as unknown parameters to be estimated. We introduce a framework to update the latent variables and parameters sequentially, with straightforward update rules. We will describe the main ingredients of the proposed framework in the following subsections. Hereafter, we use the shorthand notations 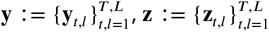, and 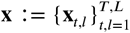.

### Preliminary assumptions

For the sake of simplicity, we assume that the constants *α*, **A**, **∑**_*w*_ and ***μ***_*x*_ are either known or can be consistently estimated from pilot trials. For example, we assume **A** to be diagonal and estimate the diagonal elements based on the magnitudes of spiking events. Further, we estimate ***μ***_*x*_ from the average firing rate and **∑**_*w*_ using the background fluorescence in the absence of spiking events. Next, we take *p*_pr_(**∑**_*x*_) to be an Inverse Wishart density:

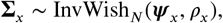

which turns out to be the conjugate prior in our model. Thus, ***ψ***_*x*_ and *ρ_x_* will be the hyper-parameters of our model. It is noteworthy that although we fix *α*, it can be updated similar to the other two hyper-parameters for better accuracy. The details of the procedure followed for hyper-parameter tuning are given in the subsection Hyper-parameter tuning.

### Decoupling via Pólya-Gamma augmentation

Direct application of VI to problems containing both discrete and continuous random variables results in intractable densities. Specifically, finding a variational distribution for **x**_*t,l*_ in our model with a standard distribution is not straightforward, due to the complicated posterior arising from co-dependent Bernoulli and Gaussian random variables. In order to overcome this difficulty, we employ Pólya-Gamma (PG) latent variables (***Pillow and Scott, 2012***; ***Polson et al., 2013***; ***Linderman et al., 2016***). We observe from ***Equation 4*** that the posterior density, *p*(**x**|**z**, **D**, **∑**_*x*_) is conditionally independent in *t,l* with:

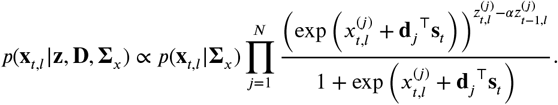

Thus, upon careful inspection we see that this density has the desired form for the PG augmentation scheme (***Polson et al., 2013***). Accordingly, we introduce a set of auxiliary PG-distributed i.i.d. latent random variables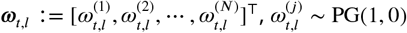 for 1 ≤ *j* ≤ *N*, 1 ≤ *t* ≤ *T* and 1 ≤ *l* ≤ *L*, to derive the complete data log-likelihood:

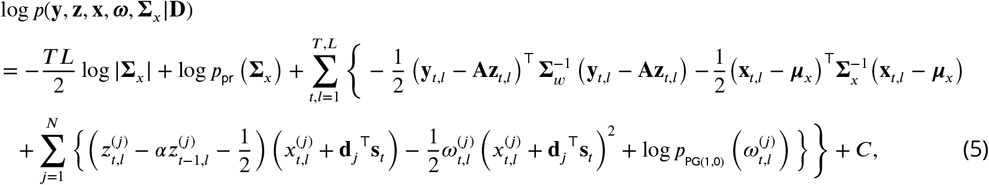

where 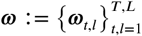 and *C* accounts for terms not depending on **y**, **z**, **x**, **ω**, **∑**_*x*_ and **D**. The complete data log-likelihood is notably *quadratic* in **z**_*t,l*_, which as we show later admits efficient estimation procedures with favorable scaling in *T*.

### Deriving the optimal variational densities

In this section, we will outline the procedure of applying VI to the latent variables 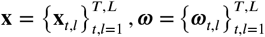 and **∑**_*x*_, assuming that the parameter estimates 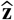 and 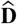 of the previous iteration are available. The methods that we propose to update the parameters 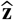 and 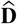 subsequently, will be discussed in the next section.

The objective of variational inference is to posit a family of approximate densities *Q* over the latent variables, and to find the member of that family that minimizes the Kullback-Leibler (KL) divergence to the exact posterior:

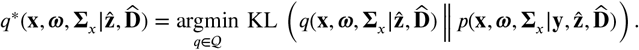

However, evaluating the KL divergence is intractable, and it has been shown (***Blei et al., 2017***) that an equivalent result to this minimization can be obtained by maximizing the alternative objective function, called the evidence lower bound (ELBO):

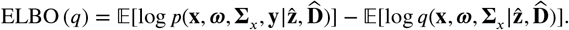

Further, we assume Q to be a mean-field variational family (***Blei et al., 2017***), resulting in the overall variational density of the form:

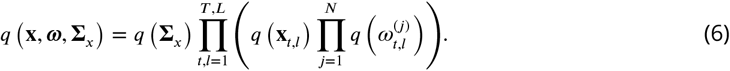

Under the mean field assumptions, the maximization of the ELBO can be derived using the optimization algorithm ‘Coordinate Ascent Variational Inference’ (CAVI) (***Bishop, 2006***; ***Blei et al., 2017***). Accordingly, we see that the optimal variational densities in ***Equation 6*** take the forms:

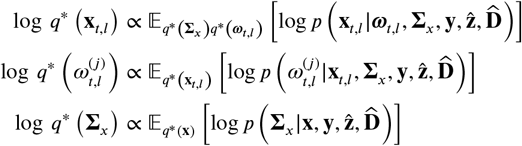

Upon evaluation of these expectations, we derive the optimal variational distributions as:

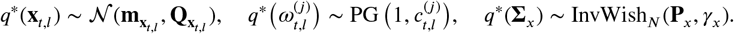

whose parameters 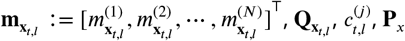, and *γ_x_* can be updated given parameter estimates 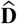 and 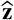:

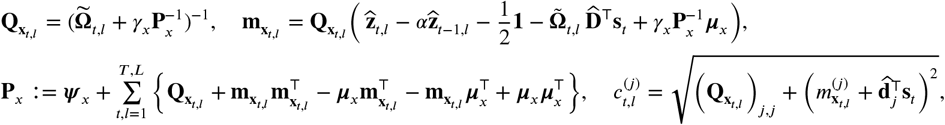

and *γ*_x_ := *ρ*_*x*_ + *T L*, with 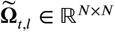 denoting a diagonal matrix with entries 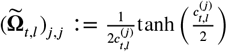 and **1** ∈ **ℝ**^*N*^ denoting the vector of all ones.

### Low-complexity parameter updates

Note that even though **z** is composed of the latent processes **z**_*t,l*_, we do not use VI for its inference, and instead consider it as an unknown parameter. This choice is due to the temporal dependencies arising from the underlying state-space model in ***Equation 4***, which hinders a proper assignment of variational densities under the mean field assumption. We thus seek to estimate both **z** and **D** using the updated variational density *q*\*(**x**, ***ω***, **∑**_*x*_).

First, note that the log-likelihood in ***Equation 5*** is decoupled in *l*, which admits independent updates to 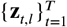 for *l* = 1,∞, *L*. As such, given an estimate 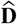, we propose to estimate 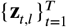 as:

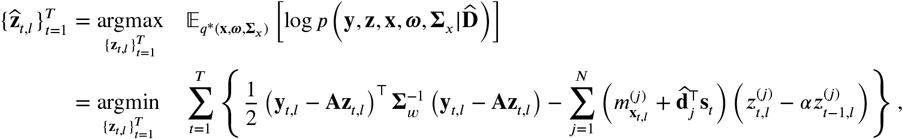

under the constraints 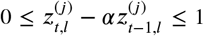, for *t* = 1,∞, *T* and *j* = 1,∞, *N*. These constraints are a direct consequence of 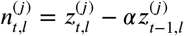 being a Bernoulli random variable with 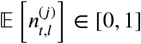. While this problem is a quadratic program and can be solved using standard techniques, it is not readily decoupled in *t*, and thus standard solvers would not scale favorably in *T*.

Instead, we consider an alternative solution that admits a low-complexity recursive solution by relaxing the constraints. To this end, we relax the constraint 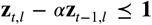 and replace the constraint 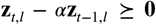 by penalty terms proportional to 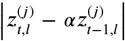. The resulting relaxed problem is thus given by:

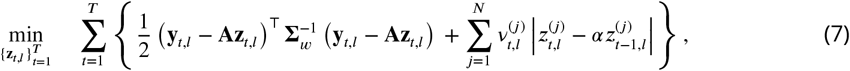

where 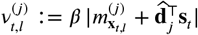 with *α* ≥ 1 being a hyper-parameter. Given that the typical spiking rates are quite low in practice, 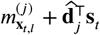 is expected to be a negative number. Thus, we have assumed that 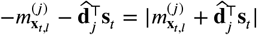.

The problem of ***Equation 7*** pertains to *compressible* state-space estimation, for which fast recursive solvers are available (***Kazemipour et al., 2018***). The solver utilizes the Iteratively Reweighted Least Squares (IRLS) (***Ba et al., 2014***) framework to transform the absolute value in the second term of the cost function into a quadratic form in **z**_*t,j*_, followed by Fixed Interval Smoothing (FIS) (***Rauch et al., 1965***) to find the minimizer. At iteration *k*, given a current estimate **z**^[*k*−1]^, the problem reduces to a Gaussian state-space estimation of the form:

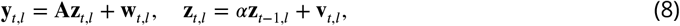

with 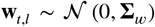 and 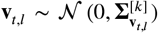, where 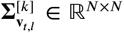 is a diagonal matrix with 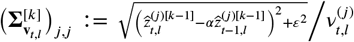, for some small constant *ε* > 0. This problem can be efficiently solved using FIS, and the iterations proceed for a total of *K* times or until a standard convergence criterion is met (***Kazemipour et al., 2018***). It is noteworthy that our proposed estimator of the calcium concentration **z**_*t,l*_ can be thought of as *soft* spike deconvolution, which naturally arises from our variational framework, as opposed to the *hard* spike deconvolution step used in two-stage estimators.

Finally, given *q*\*(**x**, ***ω***, **∑**_*x*_) and the updated 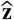, the estimate of **d**_*j*_ for *j* = 1, 2,∞, *N* can be updated in closed-form by maximizing the expected complete log-likelihood 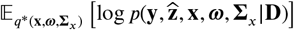:

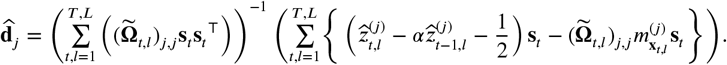

The VI procedure iterates between updating the variational densities and parameters until convergence, upon which we estimate the noise and signal covariances as:

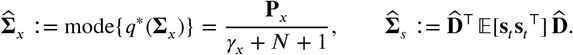

The overall combined iterative procedure is outlined in Algorithm 1. Furthermore, a MATLAB implementation of this algorithm is publicly available in ***Rupasinghe*** (***2020***).

### Model parameter settings

#### Simulation study 1

In the first simulation study, we set *α* = 0.98, *β* = 8, **A** = 0.1**I**, ***μ***_x_ = −4.5**1** and **∑**_*w*_ = 2 × 10^−4^**I** (**I** ∈ **ℝ**^8×8^ represents the identity matrix and **1** ∈ **ℝ**^8^ represents the vector of all ones), so that the SNR of simulated data was in the same range as that of experimentally-recorded data. We used a 6^th^ order autoregressive process with a mean of-1 as the stimulus (*s_t_*), and considered *M* = 2 lags of the stimulus (i.e., **s**_*t*_ = [*s_t_*, *s*_*t*-1_]^τ^) in the subsequent analysis.

#### Simulation study 2

In the second simulation study, we set *α* = 0.98, **A** = 0.1**I**, ***μ***_x_ = −4.5**1** and **∑**_*w*_ = 10^−4^**I** (**I** ∈ **ℝ**^30×30^ represents the identity matrix and **1** ∈ **ℝ**^30^ represents the vector of all ones) when generating the fluorescence traces 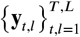, so that the SNR of the simulated data was in the same range as of real calcium imaging observations. Furthermore, we simulated the spike trains based on a Poisson process (***Smith and Brown, 2003***) using the discrete time re-scaling procedure (***Brown et al., 2002***; ***Smith and Brown, 2003***). Following the assumptions in ***Brown et al.*** (***2002***), we used an exponential link to simulate the observations:

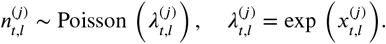

as opposed to the Bernoulli-logistic assumption in our recognition model. Then, we estimated the noise covariance 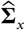 using the Algorithm 1, with a slight modification. Since there are no external stimuli, we set **s**_*t*_ = **0** and **D** = **0**. Accordingly, in Algorithm 1, we initialized 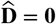 and did not perform the update on 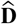 in the subsequent iterations.

**Algorithm 1.**
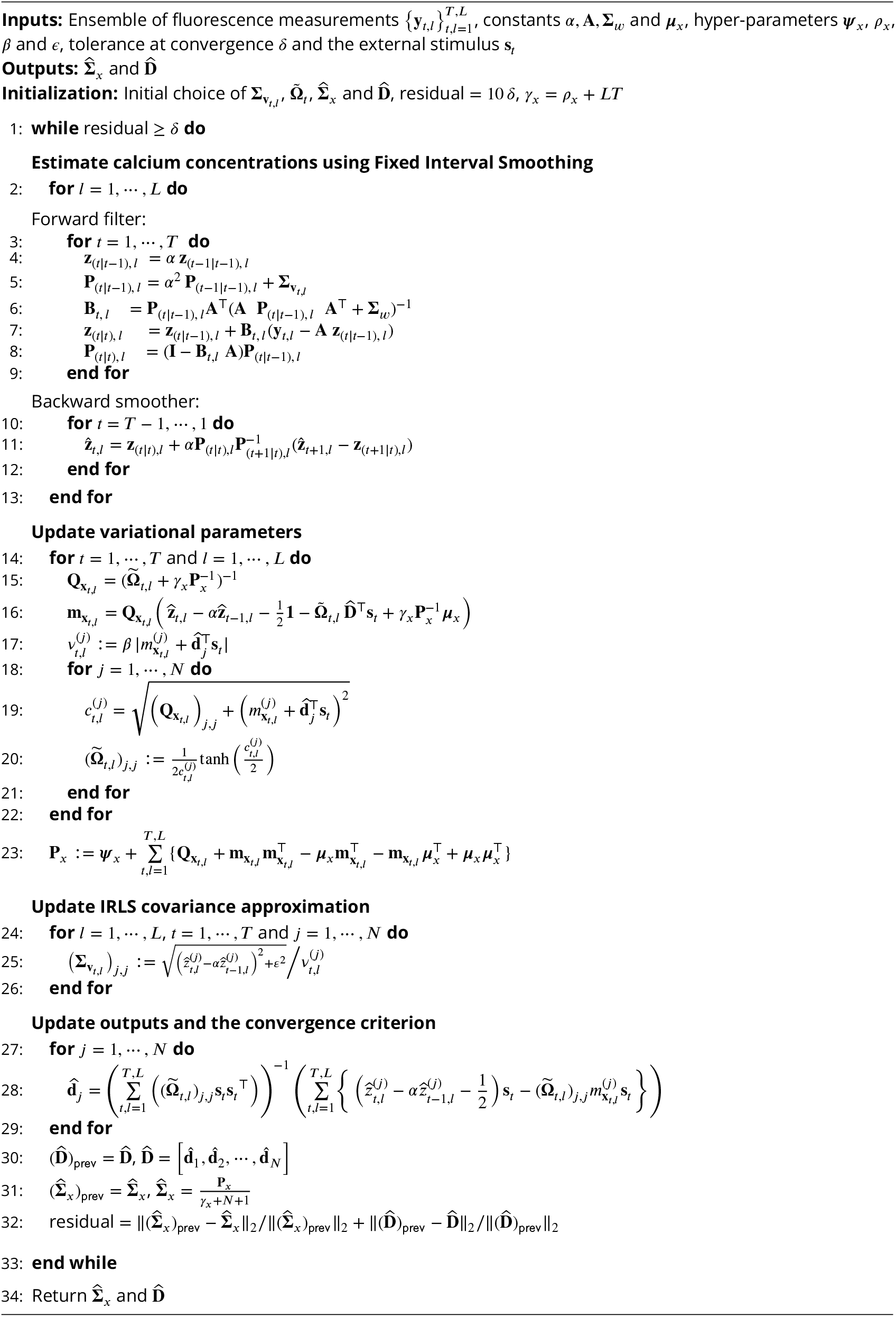
Estimation of **∑**_*x*_ and **D** through the proposed iterative procedure

#### Real data study 1

The dataset consisted of recordings from 371 excitatory neurons, from which we selected *N* = 16 neurons with high level of activity for the analysis. Each trial consisted of *T* = 3600 time frames (the sampling frequency was 30 Hz, and each trial had a duration of 120 seconds), with the presentation of a random sequence of four tones. The spiking events were very sparse and infrequent, and hence this dataset fits our model with at most one spiking event in a time frame.

We encoded the stimulus in this experiment based on the tone onsets of the four tones. Suppose that the tone onset of the *p^th^* tone (*p* =1,∞, *P*, where *P* = 4) is given by the binary sequence 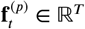. We assumed that the response at each time *t* depends only on the *R* most recent time lags of the stimulus. For each time *t*, we formulated the effective stimulus corresponding to the tone 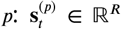, using the *R* recent lags of the tone onset sequence 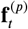 starting at *t*. Likewise, we encoded all *P* tones, and then formulated the overall effective stimulus at the *t^th^* time frame, 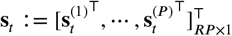. Note that the weight vector **d**_*j*_ would be *M* = *R* × *P* dimensional under this setting. Further, based on the duration of the tones and silent periods, we considered *R* = 25 time lags in this analysis.

We set *α* = 0.95 and **A** = **I** (**I** ∈ **ℝ**^16×16^ represents the identity matrix), after considering the magnitude of the spiking events in observations. Further, we estimated ***μ***_*x*_ by a linear function of average fluorescence activity. Finally, we assumed that the observation noise covariance **∑**_*w*_ is diagonal, and estimated the diagonal elements using the background fluorescence in the absence of spiking events.

#### Real data study 2

Each trial consisted of *T* = 765 frames (25.5 seconds) at a sampling frequency of 30 Hz. The layer 2/3 auditory neurons, are known to exhibit spiking rates < 5 Hz (e.g., Fig. 2-F in ***Petrus et al.*** (***2014***)), which makes the Bernoulli spiking assumption plausible for the dataset considered. Further, the auditory neurons studied here had notably low response rates (in both time and space), with only ~ 10 neurons exhibiting meaningful response. Thus, we selected *N* = 10 neurons with the highest level of activity and *L* = 10 trials for the analysis, and chose *M* = 40 lags of the stimulus in the model for the stimulus-driven condition.

We set *α* = 0.95 and **A** = 0.75**I** (**I** ∈ **R**^10×10^ represents the identity matrix), considering the magnitude of the spiking events in fluorescence observations. We used the same methods as in the first real data study to determine the optimal settings of ***μ***_*x*_ and **∑**_*w*_.

#### Real data study 3

Each experiment consisted of *L* = 5 trials of *P* = 9 different tone frequencies repeated at 4 different amplitude levels, resulting in each concatenated trial being ~ 180 second long (see ***Bowen et al.*** (***2020***) for more details). We used the same procedure as in the first real data study to encode this stimulus, setting the number of time lags to be *R* = 25. For each layer, we analyzed fluorescence observations from six experiments. In each experiment, we selected the most responsive *N* ~ 20 neurons for the subsequent analysis.

We set *α* = 0.95, **A** = **I** and used the same methods as in the previous two studies to determine the optimal settings of ***μ***_*x*_ and **∑**_*w*_ in each experiment.

### Performance evaluation

#### Simulation studies

Since the ground truth is known in simulations, we directly compared the performance of each signal and noise correlation estimate with the ground truth signal and noise correlations, respectively. Suppose the ground truth correlations are given by the matrix **X** and the estimated correlations are given by the matrix 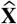. To quantify the similarity between **X** and 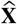, we defined the following two metrics:

##### *Normalized Mean Squared Error* (NMSE)

The NMSE computes the mean squared error of 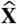 with respect to **X** using the Frobenius Norm:

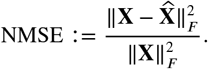

##### *Ratio between out-of-network power and in-network power* (leakage)

First, we identified the innetwork and out-of-network components from the ground truth correlation matrix **X**. Suppose that if the true correlation between the *i*^th^ neuron and the *j*^th^ neuron is non-zero, then |(**X**)_*i,j*_| > *δ_x_*, for some *δ_x_* > 0. Thus, we formed a matrix **X**^in^ that masks the in-network components, by setting (**X**^in^)_*i,j*_ = 1 if |(**X**)_*i,j*_| > *δ_x_* and (**X**^in^)_i,j_ =0 if |(**X**)_*i,j*_| ≤ *δ_x_*. Likewise, we also formed a matrix **X**^out^ that masks the out-of-network components, by setting (**X**^out^)_*i,j*_ = 1 if |(**X**)_*i,j*_| ≤ *δ_x_* and (**X**^out^)_*i,j*_ = 0 if |(**X**)_*i,j*_| > *δ_x_*. Then, using these two matrices we quantified the leakage effect of 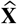 comparative to **X** by:

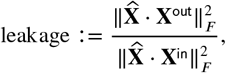

where (·) denotes element-wise multiplication.

#### Real data studies

To quantify the similarity and dissimilarity between signal and noise correlation estimates, we used a statistic based on the Tanimoto similarity metric (***Lipkus, 1999***), denoted by *T_s_*(**X**, **Y**) for two matrices **X** and **Y**. For two vectors **a** and **b** with *non-negative* entries, the Tanimoto coefficient (***Lipkus, 1999***) is defined as:

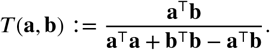

The Tanimoto similarly metric between two matrices can be defined in a similar manner, by vectorizing the matrices. Thus, we formulated a similarity metric between two correlation matrices **X** and **Y** as follows. Let **X**^+^ := max{**X**,0**I**} and **X**^−^ := max{-**X**,0**I**}, with the max{·,·} operator interpreted element-wise. Note that **X** = **X**^+^ – **X**^−^, and **X**^+^, **X**^−^ have non-negative entries. We then defined the similarity matrix by combining those of the positive and negative parts as follows:

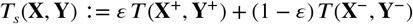

where *ε* ∈ [0.1] denotes the percentage of positive entries in **X** and **Y**. As a measure of dissimilarity, we used *T_d_*(**X**, **Y**) := 1 – *T_S_*(**X**,**Y**). The values of 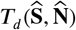 in ***Table 1*** and 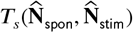 and 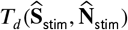 reported in ***Table 2*** were obtained based on the foregoing definitions.

To further assess the statistical significance of these results, we performed following randomized tests. To test the significance of 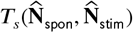, for each comparison and each algorithm, we fixed the first matrix (i.e. 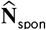) and randomly shuffled the entries of the second one (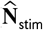 in both cases) while respecting symmetry. We repeated this procedure for 10000 trials, to derive the null distributions that represented the probabilities of chance occurrence of similarities between two random groups of neurons.

To test the significance of 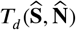 and 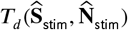, for each comparison and each algorithm, again we fixed the first matrix (i.e. signal correlations). Then, we formed the elements of the second matrix (akin to noise correlations) as follows. For each element of the second matrix, we assigned either the same element as the signal correlations (in order to model the leakage effect) or a random noise (with same variance as the elements in the noise correlation matrix) with equal probability. As before, we repeated this procedure for 10000 trials, to derive the null distributions that represent the probabilities of chance occurrence of dissimilarities between two matrices that have some leakage between them.

### Hyper-parameter tuning

The hyper-parameters that directly affect the proposed estimation are the inverse Wishart prior hyper-parameters: ***ψ***_*x*_ and *ρ_x_*. Given that *ρ_x_* appears in the form of *γ_x_* := *T L* + *ρ_x_*, we will consider ***ψ***_*x*_ and *γ_x_* as the main hyper-parameters for simplicity. Here, we propose a criterion for choosing these two hyper-parameters in a data-driven fashion, which will then be used to construct the estimates of the noise covariance matrix 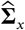 and weight matrix 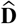. Due to the hierarchy of hidden layers in our model, an empirical Bayes approach for hyper-parameter selection using a likelihood-based performance metric is not straightforward. Hence, we propose an alternative empirical method for hyper-parameter selection as follows.

For a given choice of **ψ**_*x*_ and *γ_x_*, we estimate 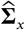 and 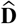 following the proposed method. Then, based on the generative model in Proposed forward model, and using the estimated values of 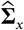 and 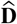, we sample an ensemble of simulated fluorescence traces 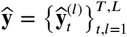, and compute the metric *d* (***ψ***_*x*_, *γ_x_*);

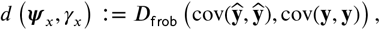

where cov(·) denotes the empirical covariance and 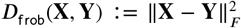. Note that *D*_frob_(**X**, **Y**) is strictly convex in **X**. Thus, minimizing *D*_frob_ (**X**, **Y**) over **X** for a given **Y** has a unique solution. Accordingly, we observe that *d*(***ψ***_*x*_, *γ_x_*) is minimized when 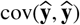 is nearest to cov(**y**, **y**). Therefore, the corresponding estimates 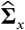 and 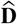 that generated 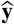, best match the second-order statistics of **y** that was generated by the true parameters **∑**_*x*_ and **D**.

The typically low spiking rate of sensory neurons observed in practice may render the estimation problem ill-posed. It is thus important to have an accurate choice of the scale matrix ***ψ***_*x*_ in the prior distribution. However, an exhaustive search for optimal tuning of ***ψ***_*x*_ is not computationally feasible, given that it has *N*(*N* + 1)/2 free variables. Thus, the main challenge here is finding the optimal choice of the scale matrix ***ψ***_*x*, opt_.

To address this challenge, we propose the following method. First, we fix ***ψ***_x,init_ = *τ***I**, where *τ* is a scalar and **I** ∈ **ℝ**^*N×N*^ is the identity matrix. Next, given ***ψ***_*x*,init_ we find the optimal choice of *γ_x_* as:

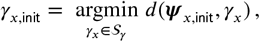

where 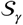. is a finite set of candidate solutions for *γ_x_* > *N* – 1. Let 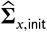 denote the noise covariance estimate corresponding to hyper-parameters (***ψ***_*x*,init_, *γ*_*x*,init_). We will next use 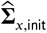 to find a suitable choice of ***ψ***_*x*_. To this end, we first fix 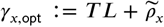, for some 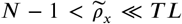. Note that by choosing 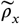 to be much smaller than *T L*, the final estimates become less sensitive to the choice of *γ_x_*. Then, we construct a candidate set *S_ψ_* for ***ψ***_*x*, opt_ by scaling 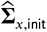 with a finite set of scalars 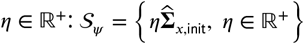. To select ***ψ***_*x*,opt_, we match it with the choice of *γ*_*x*, opt_ by solving:

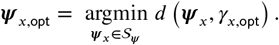

Finally, we use these hyper-parameters (***ψ***_*x*,opt_, *γ*_*x*, opt_) to obtain the estimators 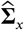 and 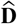 as the output of the algorithm.

### Experimental procedures

All procedures were approved by the University of Maryland Institutional Animal Care and Use Committee. Imaging experiments were performed on a P60 (for real data study 1) and P83 (for real data study 2) female F1 offspring of the CBA/CaJ strain (The Jackson Laboratory; stock #000654) crossed with transgenic C57BL/6J-Tg(Thy1-GCaMP6s)GP4.3Dkim/J mice (The Jackson Laboratory; stock #024275) (CBAxThy1), and F1 (CBAxC57). The third real data study was performed on data from P66-P93 and P166-P178 mice (see ***Bowen et al.*** (***2020***) for more details). We used the F1 generation of the crossed mice because they have good hearing into adulthood (***Frisina et al., 2011***).

We performed cranial window implantation and 2-photon imaging as previously described in ***Francis et al.*** (***2018***); ***Liu et al.*** (***2019***); ***Bowen et al.*** (***2019***). Briefly, we implanted a cranial window of 3 mm in diameter over the left auditory cortex. We used a scanning microscope (Bergamo II series, B248, Thorlabs) coupled to InsightX3 laser (Spectra-physics) (study 1) or pulsed femtosecond Ti:Sapphire 2-photon laser with dispersion compensation (Vision S, Coherent) (studies 2 and 3) to image GCaMP6s fluorescence from individual neurons in awake head-fixed mice with an excitation wavelengths of *λ* = 920 nm and *λ* = 940 nm, respectively. The microscope was controlled by ThorImageLS software. The size of the field of view was 370 × 370 *μ*m. Imaging frames of 512 × 512 pixels (pixel size 0.72 *μ*m) were acquired at 30 Hz by bidirectional scanning of an 8 kHz resonant scanner. The imaging depth was around 200 *μ*m below pia. A circular ROI was manually drawn over the cell body to extract fluorescence traces from individual cells.

#### Stimuli for real data study 1

During imaging experiments, we presented 4 tones (4, 8, 16 and 32 kHz) at 70 dB SPL. The tones were 2 s in duration with an inter-trial silence of 4 s. For the sequence of tones, we first generated a randomized sequence that consisted of 5 repeats for each tone (20 tones in total) and then the same sequence was repeated for 10 trials.

#### Stimuli for real data study 2

During imaging experiments, we presented 97 repetitions of a 75 dB SPL 100 ms broadband noise (4-48 kHz; 8 s inter-stimulus intervals). Spontaneous neuronal activity was collected from activity during 113 randomly interleaved periods of silence (8.1 s) between 1 s long noise presentations.

#### Stimuli for real data study 3

During imaging experiments, sounds were played at four sound levels (20, 40, 60, and 80 dB SPL). Auditory stimuli consisted of sinusoidal amplitude-modulated (SAM) tones (20 Hz modulation, cosine phase), ranging from 3-48 kHz. The frequency resolution was 2 tones/octave (0.5 octave spacing) and each of these tonal stimuli was 1 s long, repeated five times with a 4—6 s inter-stimulus interval (see ***Bowen et al.*** (***2020***) for details).

## Acknowledgments

This work is supported in part by the National Science Foundation Award No. 1807216 (BB) and the National Institutes of Health Award No. 1U19NS107464 (BB & POK). The authors would like to thank Daniel E. Winkowski for collecting the data in ***Bowen et al.*** (***2019***) that was also used in this work.

## Appendix 1

### Relationship to existing definitions of Signal and Noise correlations

Recall that the conventional definitions of signal and noise covariance of spiking activity between the *i*^th^ and *j*^th^ neuron are (***Lyamzin et al., 2015***):

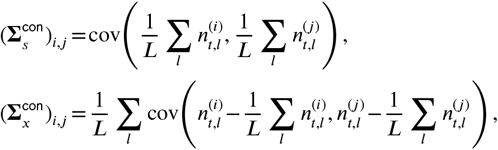

where cov 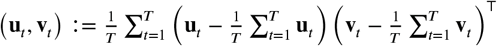, is the empirical covariance. The correlations, are then derived by the standard normalization:

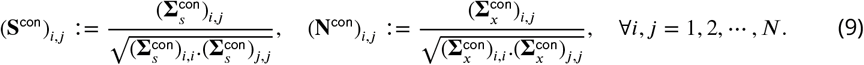

Suppose that the spiking events follow the forward model:

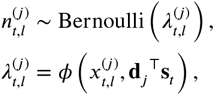

where *ϕ* : ℝ^2^ → [0.1] is a differentiable non-linear mapping. We assume **x**_*t,l*_ and **s**_*t*_, to be independent. Without loss of generality, let 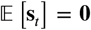 and 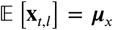. Further, we define the notation **X**_*t*_, ≈ ***Y***_*t*_, to denote almost sure equivalence, i.e., 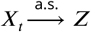 and 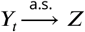 for some random variable *Z*.

First, let us consider (**S**^con^)_*i,j*_. Noting that 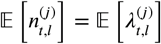 and 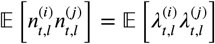, we conclude as *T* → ∞:

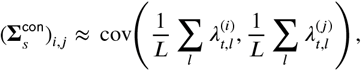

from the law of large numbers. Then, if we consider the Taylor series expansion of 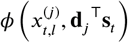,) around the mean 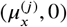, we get:

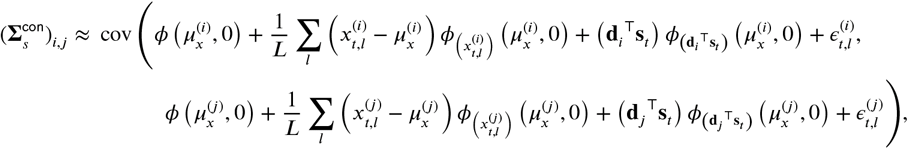

where 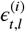 and 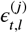 represent the higher order terms. Then, as *L* → ∞, we get:

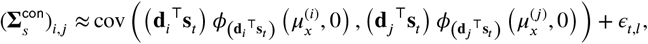

since 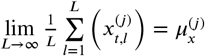 by the Law of Large numbers. Thus, we see that:

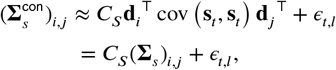

where *C_S_* is a constant and *ϵ_t,l_* is typically small if the latent process **x**_*t,l*_ and the stimulus **s**_*t*_, are concentrated around their means. Then, the signal correlations are obtained by normalization of the signal covariance as in ***Equation 9***, through which the scaling factor *C_s_* cancels and we get:

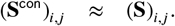

Thus, as *T,L* → ∞, we see that **S** is indeed the signal correlation matrix that is aimed to be approximated by the conventional definitions.

Next, let us consider (**N**^con^)_*i,j*_. Similar to foregoing analysis of the signal covariance, as *T* → ∞ we get:

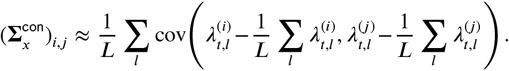

Then, from a Taylor series expansion, we get:

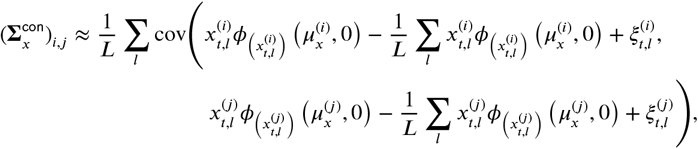

where 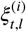 and 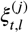 represent the higher order terms. Then, as *L* → ∞:

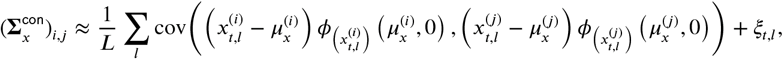

from the law of large numbers. Accordingly, we see that:

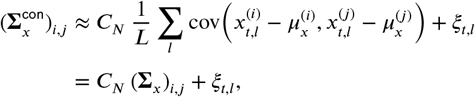

where *C_N_* is a constant and *ξ_t,l_* is typically small if the latent process **x**_*t,l*_ and the stimulus **s**_*t*_ are concentrated around their means. Then, the noise correlations are derived by normalization of the noise covariance given in ***Equation 9***. This cancels out the scaling factor *C_N_*, and we get:

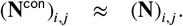

Thus, we similarly conclude that as *T,L* → ∞, the conventional definition of noise correlation **N**^con^ indeed aims to approximate **N**.

As a numerical illustration, we demonstrated in ***Figure 2-Figure Supplement 2*** that the conventional definitions of the correlations indeed approximate our proposed definitions, but require much larger number of trials to be accurate. More specifically, in order to achieve comparable performance to our method using *L* = 20 trials, the conventional correlation estimates require *L* = 1000 trials.

## Appendix 2

### Proof of Theorem 1

In what follows, we present a comprehensive proof of Theorem 1. Recall the following key assumptions:

#### Assumption (1).

We assume a scalar time-varying external stimulus (i.e. **s**_*t*_ = *s_t’_*, and hence **d**_*j*_ = *d_j_*, **d** = [*d*_1_, *d*_2_, ∞, *d_N_*]^*T*^). Furthermore, we set the observation noise covariance to be 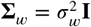, for notational convenience.

#### Assumption (2)

We derive the performance bounds in the regime where *T* and *L* are large, and thus do not impose any prior distribution on the correlations (i.e., *p*_pr_(**∑**_*x*_) ∞ 1), which are otherwise needed to mitigate overfitting (see Methods and Materials).

#### Assumption (3).

We assume the latent noise process and stimulus to be slowly varying signals, and thus adopt a piece-wise constant model in which these processes are constant within consecutive windows of length *W* (i.e., **x**_*t,l*_ = **x***_W_k_l_* and *s_t_* = *s_W_k’__*, for (*k*-1)*W* + 1 ≤ *t* < *kW* and *k* = 1, ∞, *K* with *W_k_* = (*k*-1)*W* + 1 and *KW* = *T*) for our theoretical analysis, as is usually done in spike count calculations for conventional noise correlation estimates.

#### Proof of Theorem 1.

First, recall the proposed forward model (see Methods and Materials) under Assumption (1)-(3):

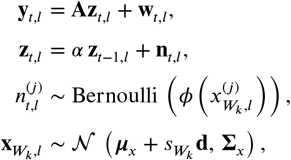

where 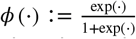, is the logistic function. Note that we have re-defined the latent process **x**_*t,l*_ by absorbing the stimulus activity *s_t_***d** to the mean of **x**_*t,l*_ for notational convenience, without loss of generality. Hereafter, we also assume that **A** = **I** without loss of generality. For a truncation level ***B*** (to be specified later), consider the event 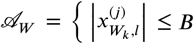 and 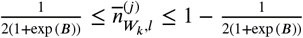 for *j* = 1,∞, *N, k* = 1,∞, *K* and *l* = 1, ∞ *L*}, such that 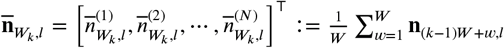. First, we derive convenient forms of the maximum likelihood estimators via the Laplace’s approximations and asymptotic expansions (***Wong, 2001***) through the following lemma:

#### Lemma 1.

*Conditioned on event 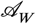, the maximum likelihood estimators of the stimulus kernel of the *j^th^* neuron and the noise covariance between the *i^th^* and *j^th^* neurons take the forms*:

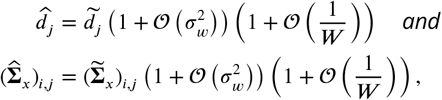

*where*

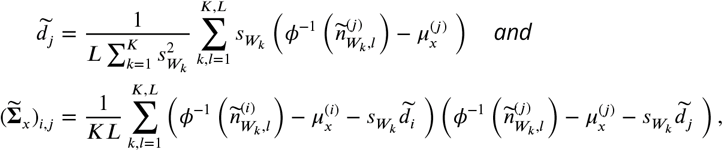

*with* 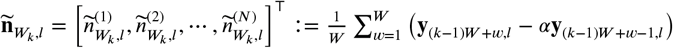 *and* 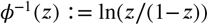.

#### Proof of Lemma 1.

First, maximizing the data likelihood, we derive the estimators:

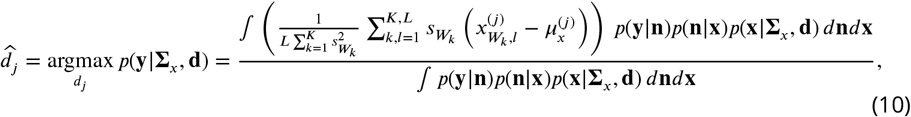

and

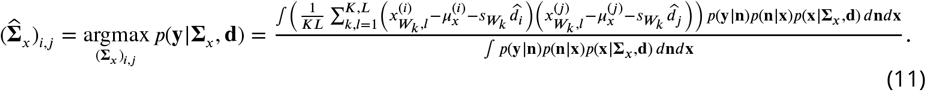

where *W_k_* = (*k* - 1)*W* + 1. Then, we simplify these integrals based on the saddle point method of asymptotic expansions (***Wong, 2001***). To that end, first consider the numerator of ***Equation 10*** denoted by 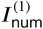. First, we evaluate the integration in 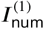 with respect to the variable **n**. To that end, note:

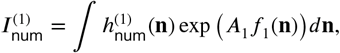

where 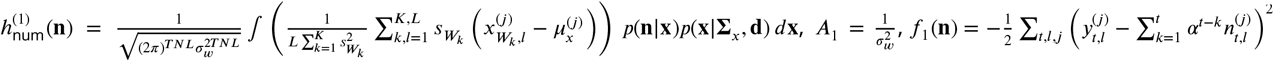 and *d***n** is shorthand notation for the product measure of the discrete random vector **n**. Observing that 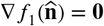 for 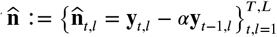, using the method of asymptotic expansions, 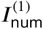, can be evaluated as:

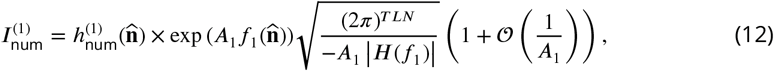

where the determinant of the Hessian matrix |*H*(*f*_1_)|, is a negative function of *α*. Note that the covariance of this Gaussian integral [-(*H*(*f*_1_)^−1^) is a function of *α* ∈ (0.1), and hence is bounded. Thus, all higher order error terms in ***Equation 12*** are also bounded, as higher order moments of Gaussian distributions are functions of the covariance.

Next, we simplify the integral 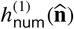 in ***Equation 12*** using a similar procedure. We have:

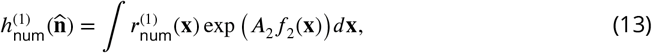

where 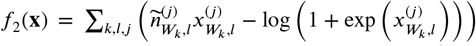 with 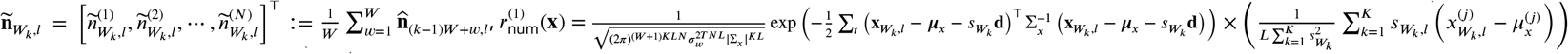 and *A*_2_ = *W*. Then, we note that the gradient of 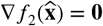 for 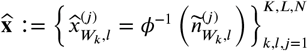, where *ϕ*^−1^(*z*) := logit(*z*) = l⊓(*z*/(1 - *z*)). Accordingly, by re-applying the saddle point method of asymptotic expansions, we evaluate the integral in ***Equation 13*** as:

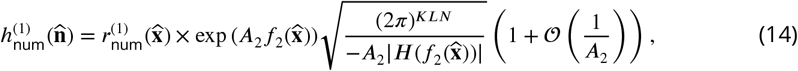

where the determinant of the Hessian,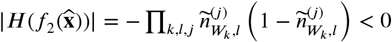 when conditioned on event 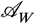. The higher order terms in ***Equation 14*** will be bounded if the covariance of the saddle point approximation 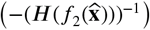 is bounded, which we ensure by conditioning on event 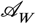. This completes the evaluation of 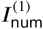.

Following the same sequence of arguments, we evaluate the denominator of ***Equation 10*** denoted by 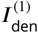. Accordingly, we derive:

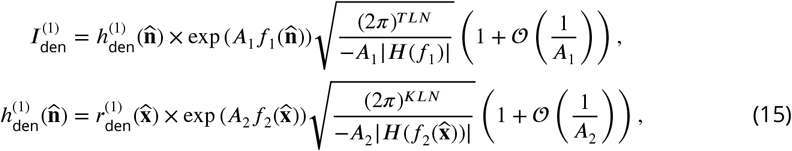

where 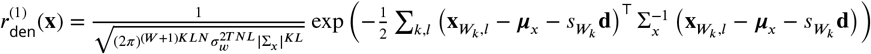. Finally, by combining ***Equation 12, Equation 14*** and ***Equation 15***, the maximum likelihood estimator in ***Equation 10*** takes the form:

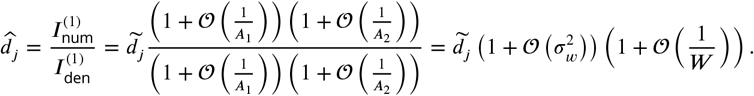

Further, following the same sequence of reasoning, simplifying the numerator 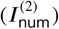 and denominator 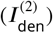 of ***Equation 11*** yields:

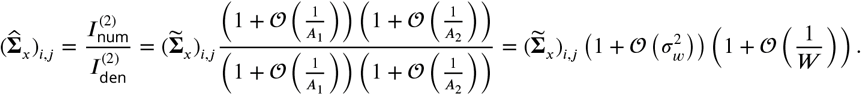

This concludes the proof of Lemma 1.

Given that *ϕ*^−2^(*z*) is unbounded for *z* = 0 or *z* = 1, we consider another truncation: 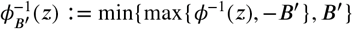, where *B*’ = 2log (2exp (*B*) + 1). This choice of *B*’ guarantees that over 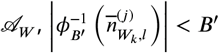 for all *j* = 1,∞, *N, k* =1,∞, *K* and *l* = 1,∞, *L:* and thus 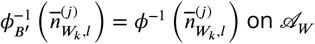.

From Lemma 1, the bias and variance of the maximum likelihood estimators, 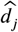 and 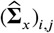 are upper-bounded, if those of 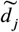 and 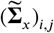 are bounded:

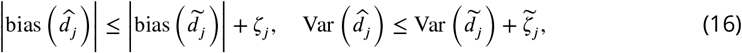

and

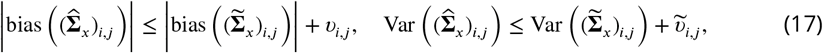

where 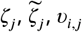 and 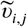 represent terms that are 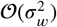 or 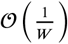. (Thus, we seek to derive the performance bounds of 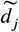 and 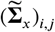.

Bounding the bias of 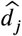

Let us first consider 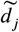. Note that:

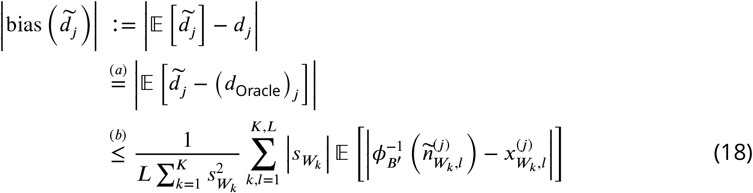

where (*a*) holds since the Oracle estimator, 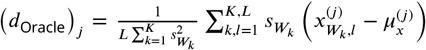 (i.e., observing **x**_*t,l*_ directly) is unbiased and (*b*) follows through the application of Jensen’s inequality and triangle inequality. To simplify this bound, the triangle inequality yields:

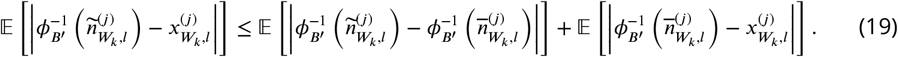

Then, to bound each of these terms, we establish a piece-wise linear Lipschitz-type bound on 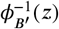. First, consider the first term 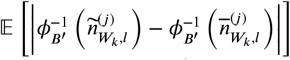. We seek to upper-bound this expectation by bounding 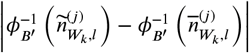 via the following technical lemma:.

#### Lemma 2.

*Conditioned on event 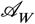, the following bound holds for all j* = 1, ∞, *N, k* =1,∞,*K and l* = 1, ∞, *L*:

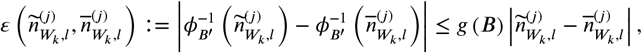

where

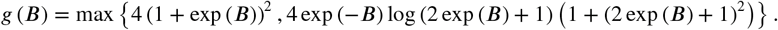

#### Proof of Lemma 2.

First, consider the case 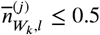. We bound the function 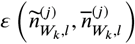 in a piece-wise fashion as follows. Note that 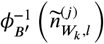 is convex for 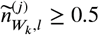 and concave for 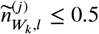. Thus, it immediately follows that for, 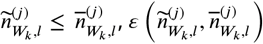 is convex and hence:

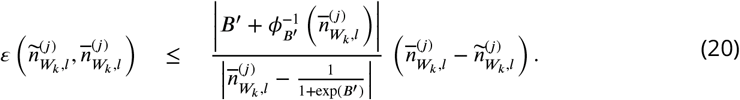

Furthermore, for 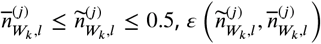 is concave, and hence is bounded by the tangent at 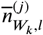:

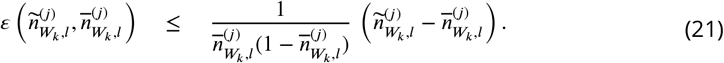

Finally, for the case of 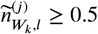, consider the line,

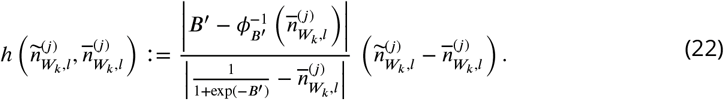

From the convexity of 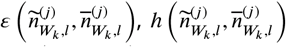 upper bounds 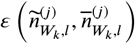 for 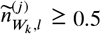, since 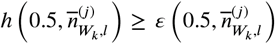 for 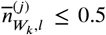. Combining the piece-wise bounds in ***Equation 20, Equation 21*** and ***Equation 22***, we conclude that for 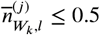:

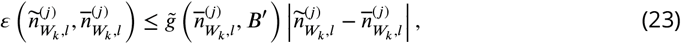

where

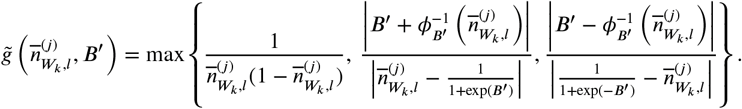

Due to the symmetry of 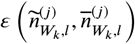, the same bound in ***Equation 23*** can be established for 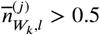 as well.

Then, using 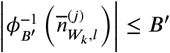 and conditioning on event 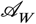, we simplify this bound as:

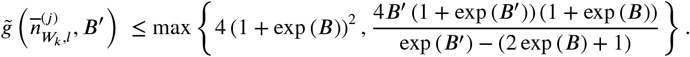

Finally, based on the fact that *B*’ = 2log(2exp(*B*) + 1), the latter is further upper bounded as:

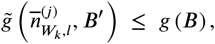

where

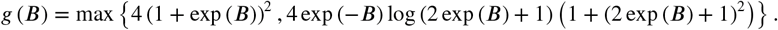

This concludes the proof of Lemma 2.

Following Lemma 2, by conditioning on the event 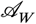 we have:

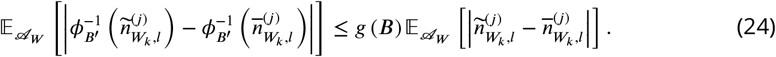

Then, we note that:

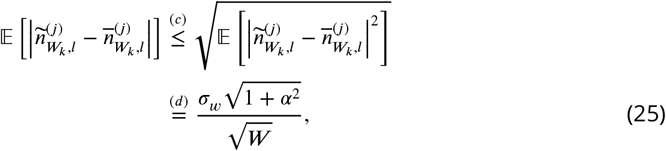

where in (*c*) we have used the Cauchy-Schwarz inequality, and in (*d*) we have used the fact that the observation noise across the *W* time instances is i.i.d. and white. From the bounds in ***Equation 24*** and ***Equation 25***, we conclude that the first expectation in ***Equation 19***, conditioned on event 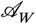 is bounded as:

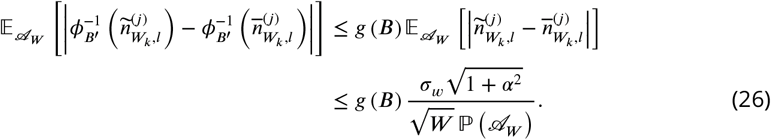

The foregoing sequence of reasoning similarly follows for 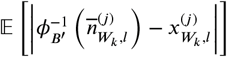, since 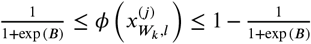 for *k* =1,∞, *K, l* = 1,∞, *L* and *j* = 1,∞, *N* (as a consequence of 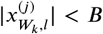 for *k* =1,∞, *K, l* = 1,∞, *L* and *j* = 1,∞, *N*, conditioned on 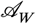). Accordingly,we derive the upper bound on the second term in ***Equation 19***, conditioned on event 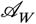:

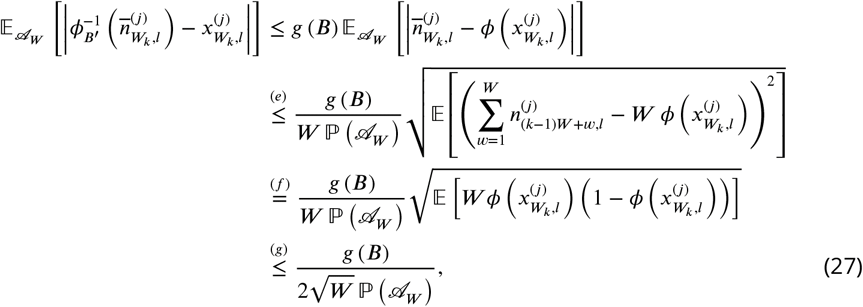

where (*e*) follows from the application of Jensen’s inequality, (*f*) follows from the formul) for the variance of a Binomial random variable, and (*g*) follows from the inequality 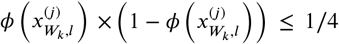, for 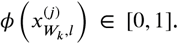 Combining the results in ***Equation 26*** and ***Equation 27***, the overall expectation in ***Equation 19***, conditioned on the event 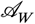 is upper- bounded by:

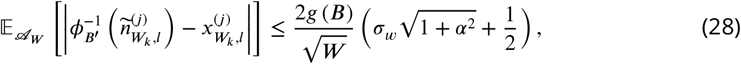

where we have lower bounded the probability of the event 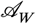 by 1/2 (that is, 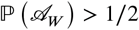). Thus, from ***Equation 18*** and ***Equation 28*** we derive:

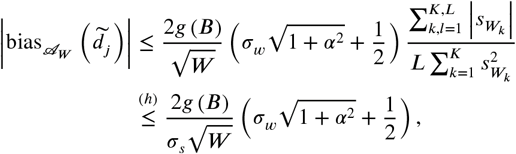

where in (*h*) we have used the Cauchy-Schwarz inequality 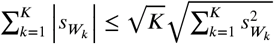 while defining 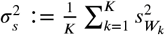.

Then, for ***B*** ≥ 2.5, we have *g* (***B***) = 4(1 + exp(***B***))^2^ and *B*’ = 2log(2exp(*B*) + 1) ≤ 3*B*. Let 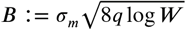 for some 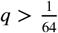. Further, for some *ϵ* < 1/2, suppose that:

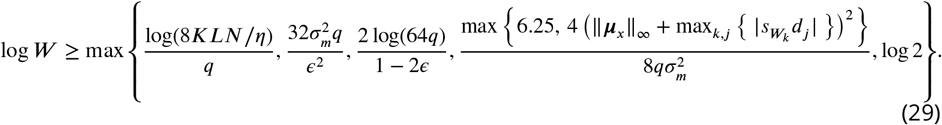

Under these conditions,

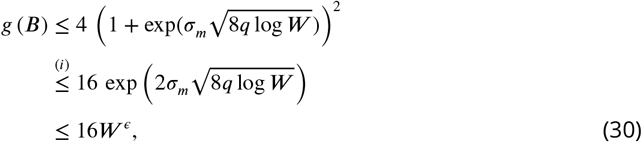

where in (*i*) we have used the fact that *e^x^* ≥ 1 for *x* ≥ 0. Thus, under the conditions in ***Equation 29***, we have:

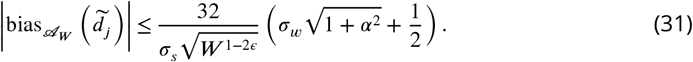

Finally, from ***Equation 16*** and ***Equation 31***, we conclude that:

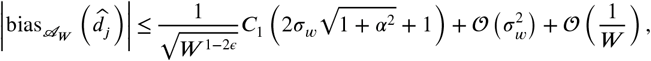

where 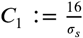.

Bounding the variance of 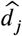

Next, we prove the upper bound on the variance of the maximum likelihood estimator, 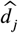. To that end, we upper-bound the variance of 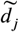. First, using the Cauchy-Schwarz inequality, we have:

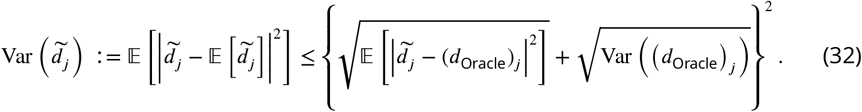

Then, we upper-bound the conditional second moment of 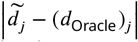 using the same techniques as we used in bounding the first moment. Accordingly, we get:

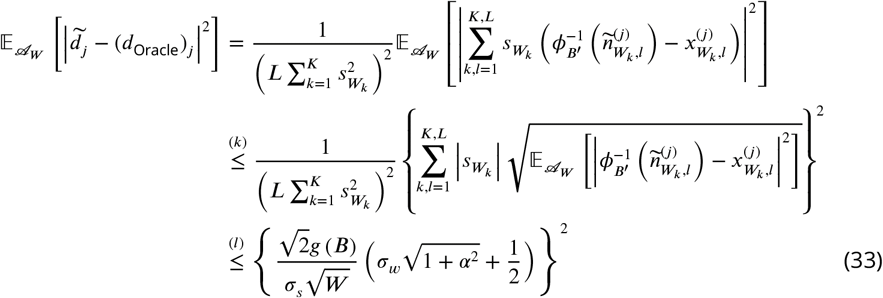

where in (*k*) we have used the Cauchy-Schwarz inequality and (*l*) follows from 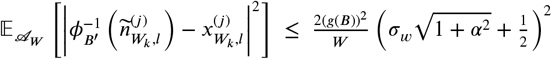, which can be proven by the same techniques as before.

Next, we note that the variance of the Oracle estimator (*d*_Oracle_)_*j*_:

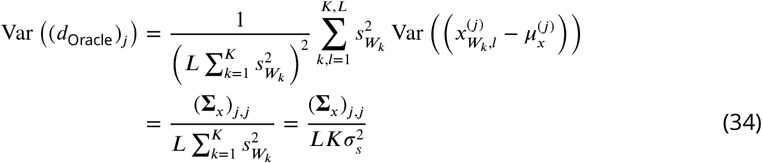

Combining ***Equation 32, Equation 33*** and ***Equation 34***, we can upper-bound the conditional variance of 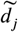 as:, following ***Equation 32***:

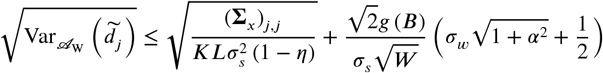

Then, following ***Equation 16***, under the conditions for *W* in ***Equation 29***, we conclude the proof of the conditional variance of 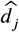:

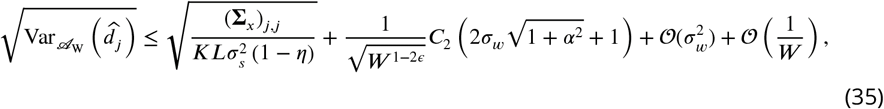

where 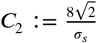.

Bounding the bias of 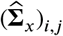.

Next, following the foregoing techniques, we upper-bound the bias and variance of the noise covariance estimator 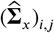. To that end, we first note:

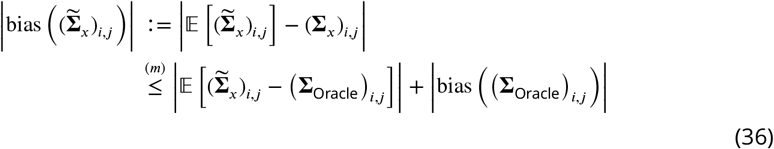

where (*m*) follows from the triangle inequality, with the Oracle noise covariance estimator (i.e., observing **x**_*t,l*_ directly) being defined as: 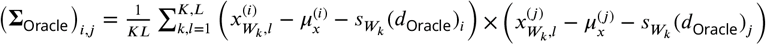. Then, to simplify the first term in ***Equation 36***, we use similar techniques as before. Accordingly,

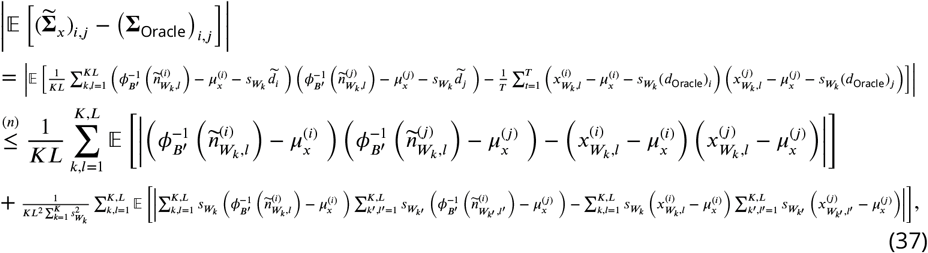

where (*n*) follows through the application of Jensen’s inequality and triangle inequality. Next, we have:

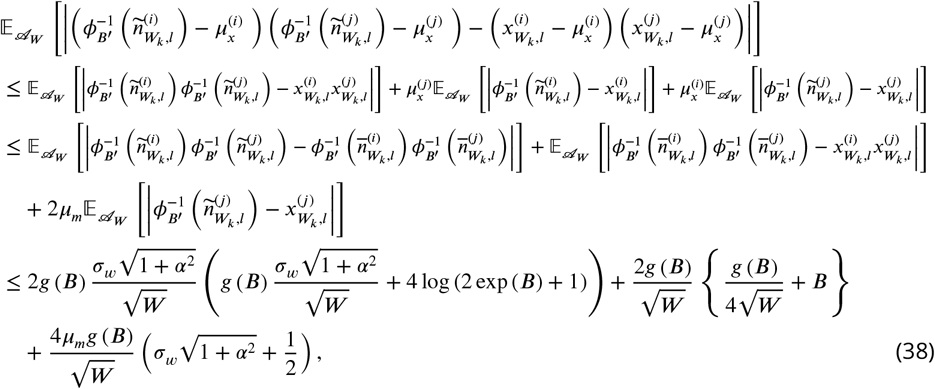

where *μ_m_* = ||***μ***_*x*_||_∞_ and we have used *B*’ = 2log(2exp(*B*) + 1). Similarly, the second term in ***Equation 37*** can be bounded as:

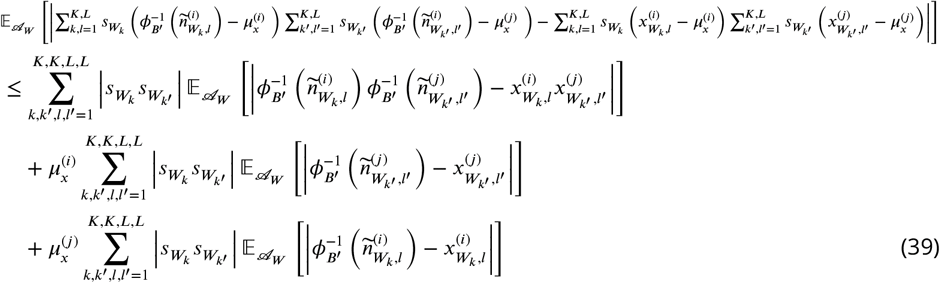

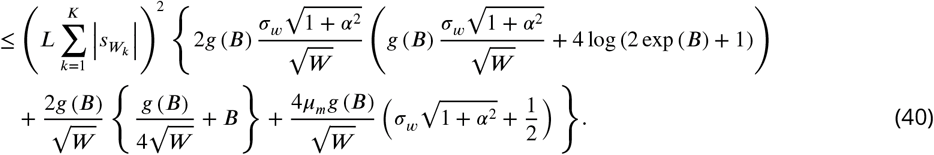

Then, by combining the bounds in ***Equation 38*** and ***Equation 40*** and using an instance of Cauchy-Schwarz inequality 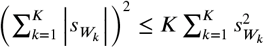, we see that the bound in ***Equation 37*** can be expressed as:

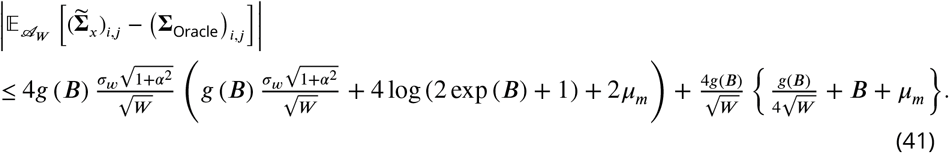

Next, we see that the oracle estimator follows an Inverse Wishart distribution, that is *KL***∑**_Oracle_ ~ Inv Wish_N_(**∑**_*x*_, *KL* - 1). Therefore, we get:

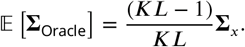

Thus, the bias of the oracle estimator is given by:

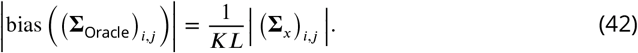

Combining the results in ***Equation 41*** and ***Equation 42***, the bias of 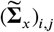 can be bounded as:

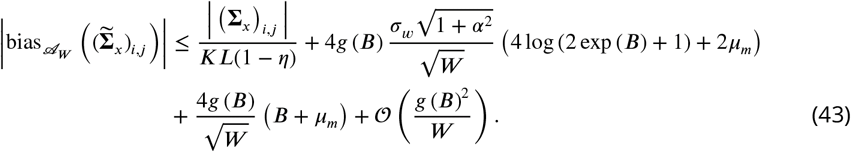

Finally, under the conditions for *W* in ***Equation 29***, the latter inequality simplifies to:

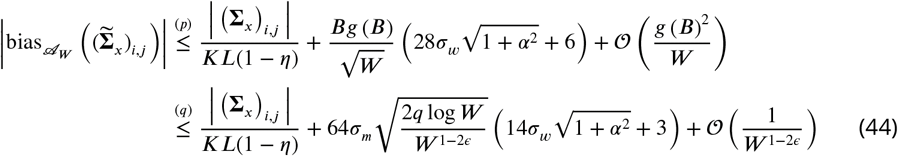

where in (*p*) we have used 2log(2exp(*B*) +1) ≤ 3*B* and *B* > 2*μ_m_* and in (*q*) we have used 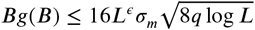, which follows from ***Equation 30***. Thus, following ***Equation 17*** we derive the bound on the bias of the maximum likelihood estimator:

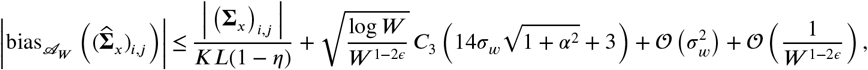

where 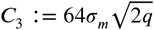.

Bounding the variance of 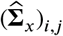

Next, we establish an upper bound on the variance of the maximum likelihood estimator of the noise covariance. To that end, we upper-bound the variance of 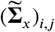. First, using the Cauchy-Schwarz inequality, we get:

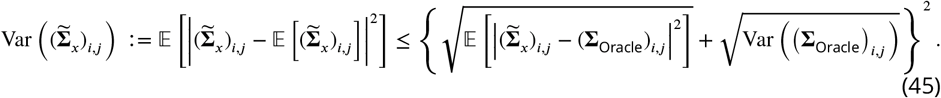

Then, we upper-bound the conditional second moment of 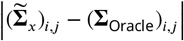 using the same techniques used in bounding its first moment. Accordingly, we derive:

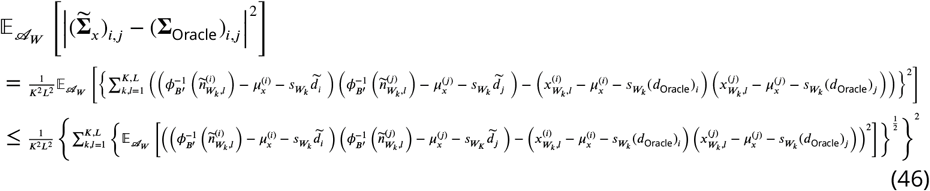

where the last bound follows from the Cauchy-Schwarz inequality. Then, we derive:

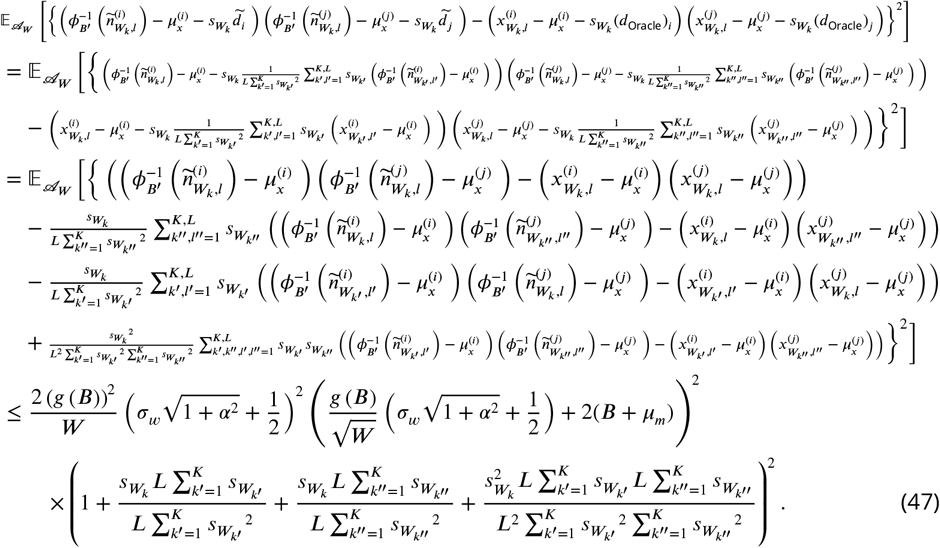

Using the final bound of ***Equation 47*** in ***Equation 46***, we get:

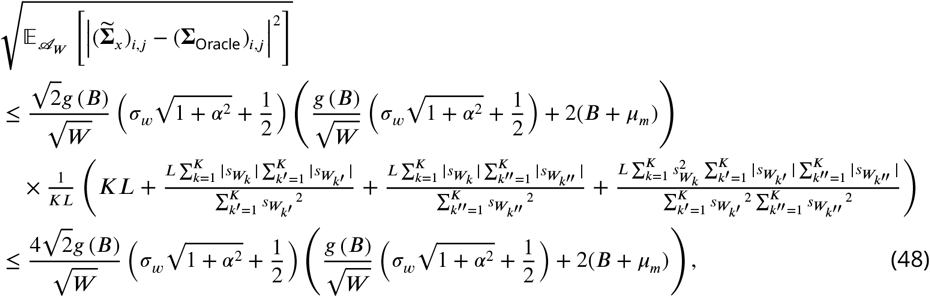

where the last inequality follows from an instance of the Cauchy-Schwarz inequality, i.e., 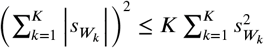.

Then, following the observation *KL***∑**_Oracle_ ~ InvWish_N_(**∑**_*x*_. *KL* - 1), we derive the variance of (∑_Oracle_)_*i, j*_:

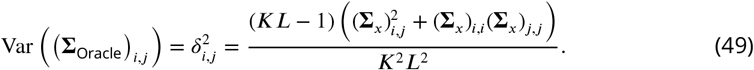

Combining ***Equation 45, Equation 48*** and ***Equation 49***, we express the upper bound on the conditional variance of 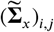 as:

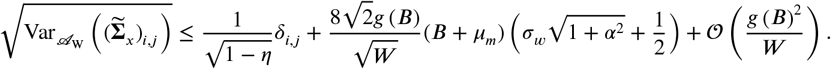

Then, following ***Equation 17*** and the conditions in ***Equation 29***, we conclude the proof of the upper bound on the conditional variance of 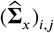:

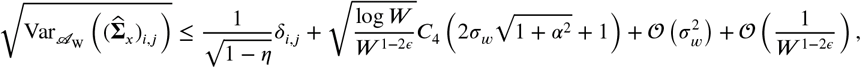

Finally, it only remains to prove that the event 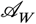 occurs with high probability for sufficiently large *W*:

#### Lemma 3.

*The probability of occurrence of the event {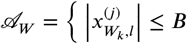 and 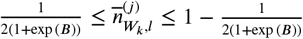 for *j* = 1,∞, *N,k* = 1,∞, *K* and *l* = 1, ∞, *L*} is upper-bounded as follows*:

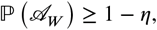

*for some constant* 0 < *η* ≤ 1/2 *satisfying the conditions of Eq. (29).*

#### Proof of Lemma 3.

First, using the union bound, we have:

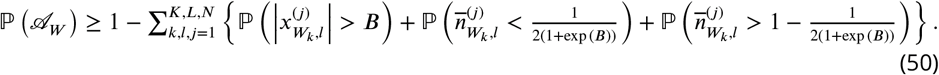

Next, we bound the probabilities on the right hand side using Chernoff’s inequality (***Boucheron et al., 2013***). First, note that:

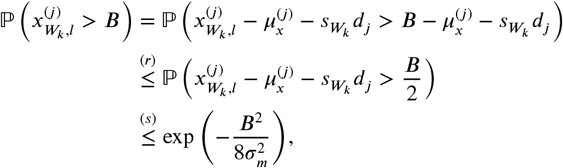

where (*r*) follows if 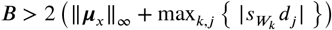 (which will hold under the conditions in ***Equation 29***) and (*s*) has been derived by applying the Chernoff’s bound on the Gaussian random variable 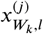. From the same reasoning we see that 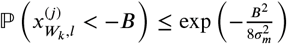. Combining these two results, we get the upper bound:

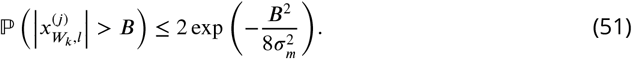

Next, note that:

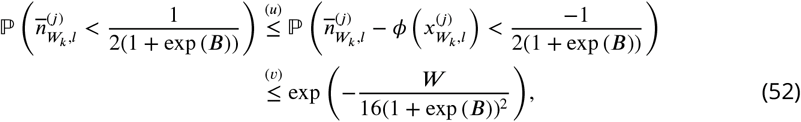

where (*u*) follows from the observation 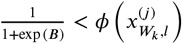 (which is a consequence of 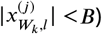. Then, we note that the zero-mean random variable 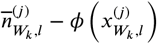 is sub-Gaussian with variance factor 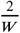. Thus, using the Chernoff’s inequality on sub-Gaussian random variables (***Boucheron et al., 2013***), we derive the upper-bound (*v*) in ***Equation 52***. In a similar fashion, based on the observation 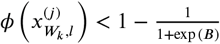, we conclude the bound:

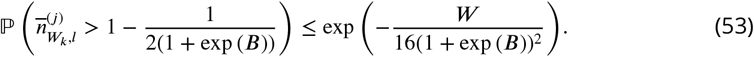

By combining the bounds in ***Equation 51, Equation 52*** and ***Equation 53***, the upper bound on 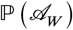 in ***Equation 50*** takes the form:

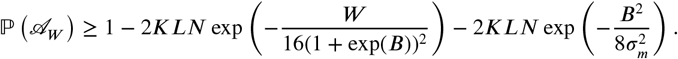

Finally, under the assumptions in ***Equation 29***, we further simplify this bound as:

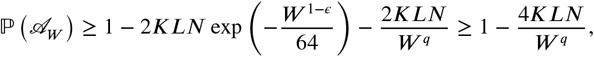

where we have used *W* ≥ 2 (which gives log *W* ≥ 2loglog *W*) and log 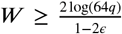 to show that 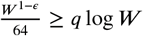. Thus, log 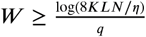 ensures that 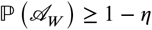 for 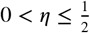.

This concludes the proof of Theorem 1.

---

**Figure 2-Figure supplement 1.**
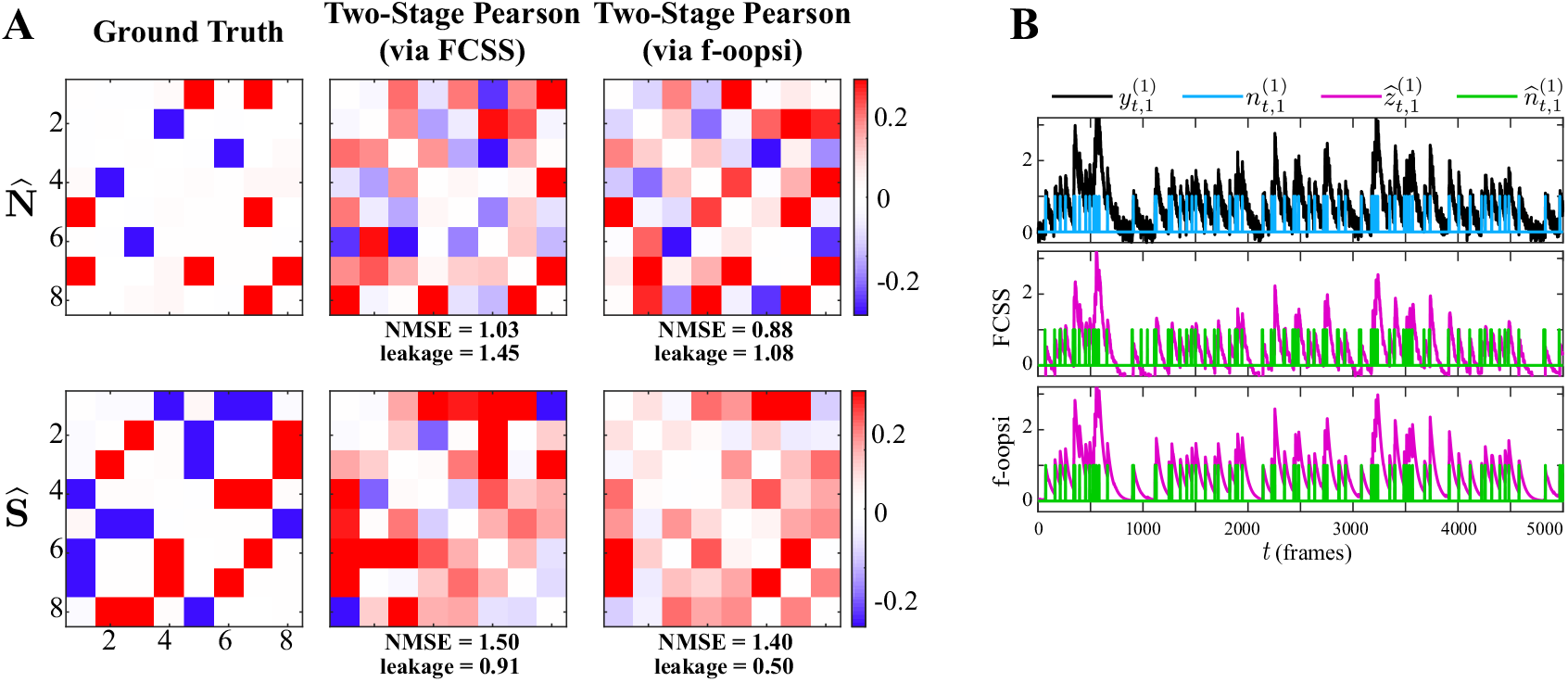
A) Noise (first row) and signal (second row) correlations corresponding to the ground truth (first column), estimated by the two-stage Pearson method using the FCSS (***Kazemipour et al., 2018***) (second column) and constrained f-oopsi (***Pnevmatikakis et al., 2016***) (third column) spike deconvolution techniques, for the simulation study in **Figure 2**. The NMSE and leakage ratios of the estimates are indicated below each panel. While the correlation estimates based on these two methods are comparable, there exist notable differences between them, as a result of the slight discrepancies in the deconvolved spikes. This demonstrates that the two-stage estimates are notably sensitive to minor differences in the estimated spikes obtained by different deconvolution techniques. In addition, both two-stage Pearson estimates fail to capture the ground truth correlations (as is also evident from the high NMSE and leakage values). B) Simulated observations (black, re-scaled for ease of visual comparison) and ground truth spikes (blue), as well as the estimated calcium concentrations (purple) and putative spikes (green) for the 1^st^ trial of neuron 1 in the simulation study of **Figure 2**, using the FCSS (***Kazemipour et al., 2018***) (second row) and constrained f-oopsi (***Pnevmatikakis et al., 2016***) (third row) spike deconvolution methods.

**Figure 2-Figure supplement 2.**
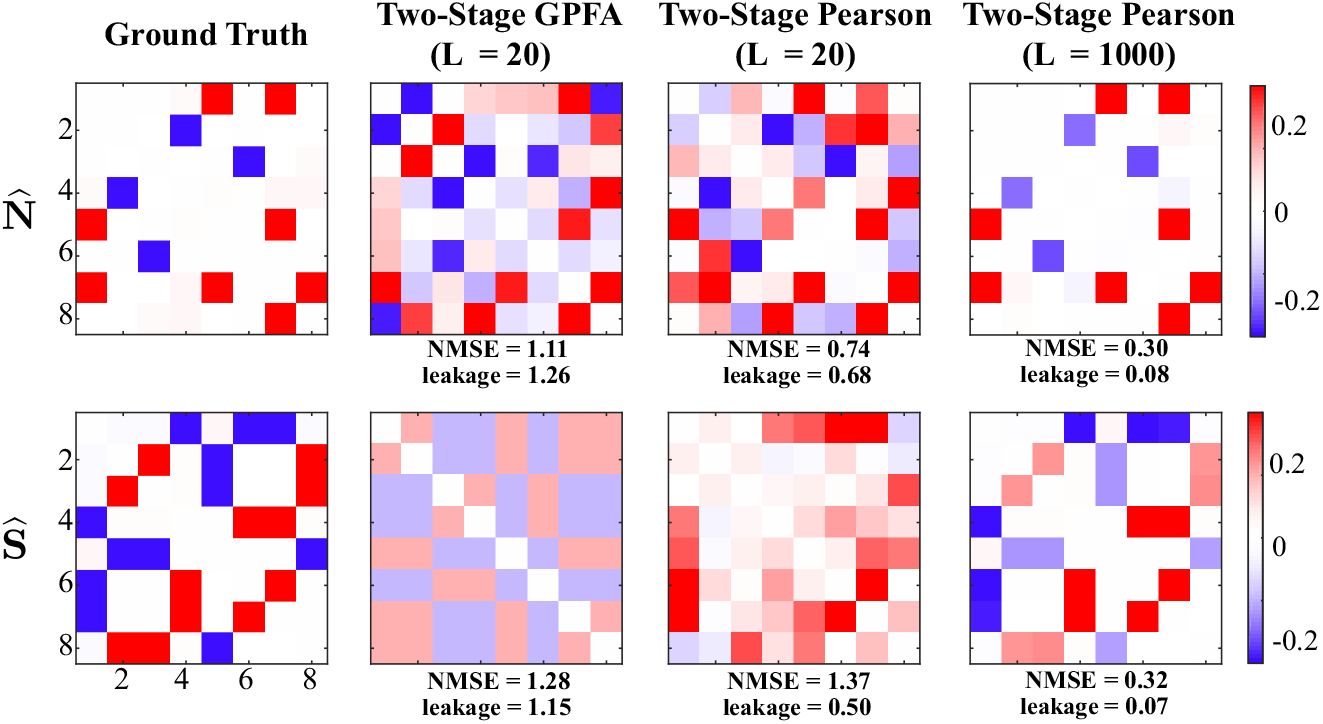
Performance of two stage estimates based on ground truth spikes. Noise (first row) and signal (second row) correlations corresponding to the ground truth (first column) are repeated from **Figure 2**. The second and third columns show the results of two-stage GPFA and two-stage Pearson methods using *L* = 20 trials, respectively. The fourth column shows the results of the two-stage Pearson method using *L* = 1000 trials. All estimates were obtained using the ground truth spikes, as opposed to extracting the spikes via a deconvolution technique. Thus, these results isolate the effect of the non-linearities involved in spike generation on the estimation performance. The NMSE and leakage ratios of the estimates are indicated below each panel. Even though the ground truth spikes are used, the NMSE and leakage ratios indicated in the second and third columns are remarkably high. This further shows that the usage of conventional definitions and GPFA estimates is not optimal for the recovery of signal and noise correlations. In accordance with our theoretical analysis in **Appendix 1**, the performance of the two-stage Pearson method significantly improves as the number of trials is increased to *L* = 1000, a number that is unrealistic in the context of typical two-photon imaging experiments. However, our proposed method shown in **Figure 2** achieves comparable performance with number of trials as low as *L* = 20. In summary, these results suggest that the two-stage methods produce highly biased estimates under limited number of trials, even if the ground truth spikes were ideally deconvolved from the two-photon data.

**Figure 2-Figure supplement 3.**
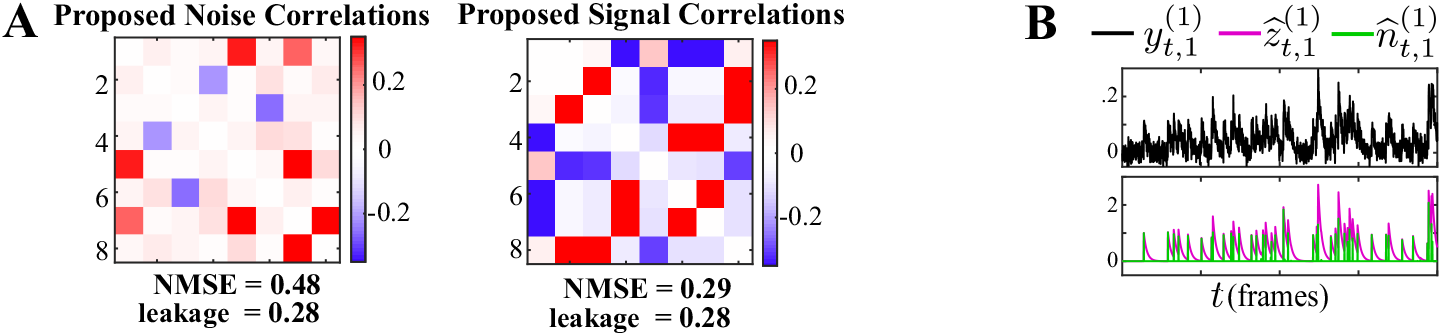
A) Proposed noise and signal correlation estimates for data simulated at lower SNR than the setting of **Figure 2** and model mismatch introduced by using a second-order autoregressive model for the calcium decay. The ground truth correlations are the same as those in **Figure 2**. The NMSE and leakage ratio are given at the bottom. B) putative spikes (green) and estimated calcium concentrations (purple). The model mismatch and lower SNR result in slight performance degradation compared to **Figure 2** (in terms of NMSE and leakage), and our method is capable of recovering the underlying correlations faithfully.

**Figure 6-Figure supplement 1.**
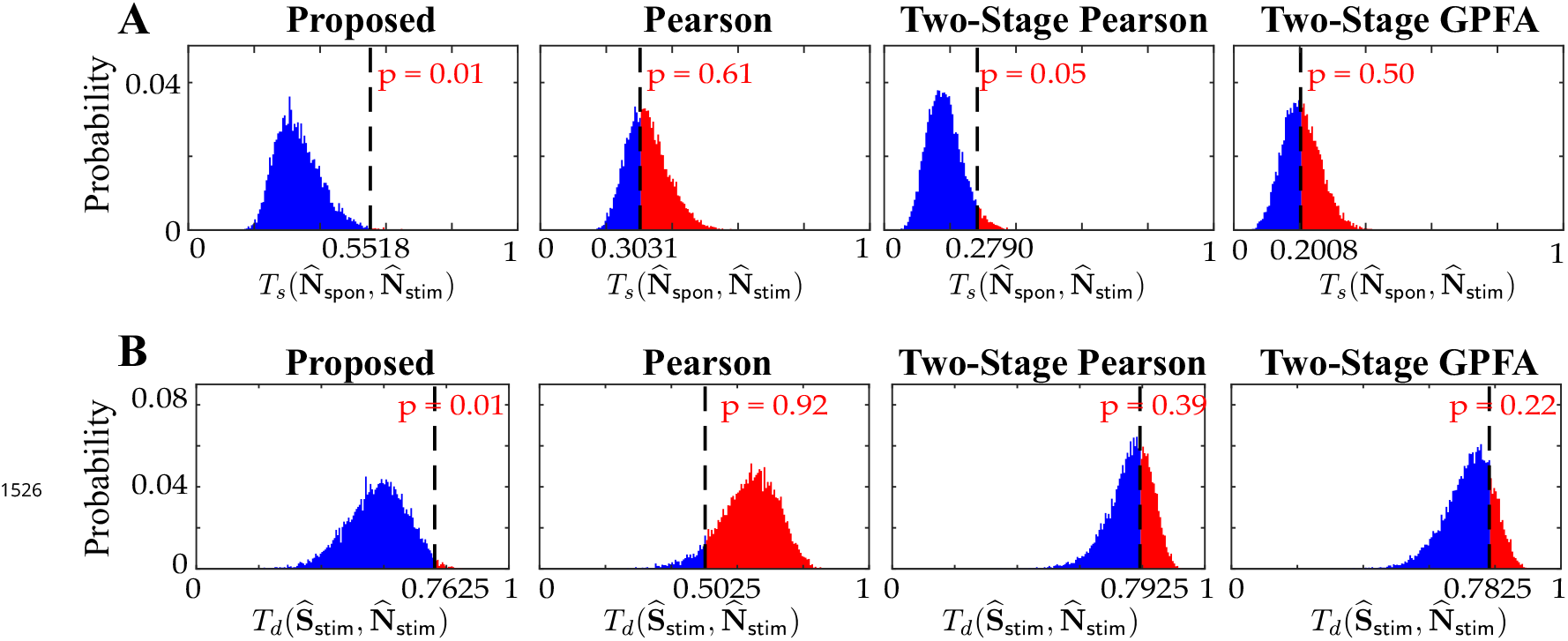
Null distributions of A) the similarities between **N**_spon_ and **N**_stim_ (top: 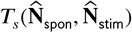) and B) the dissimilarities between 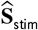 and 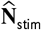 (bottom: 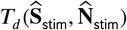), obtained by the shuffling procedure applied to the results of real data study 2 in **Figure 6**. The observed test statistic in each case is indicated by a dashed vertical line. Rows from left to right: proposed method, Pearson correlations from two-photon data, two-stage Pearson correlations and two-stage GPFA estimates. These results show that the only statistically significant outcomes (with *p* ≤ 0.05) are the similarities and dissimilarities obtained by our proposed method.

**Figure 7-Figure supplement 1.**
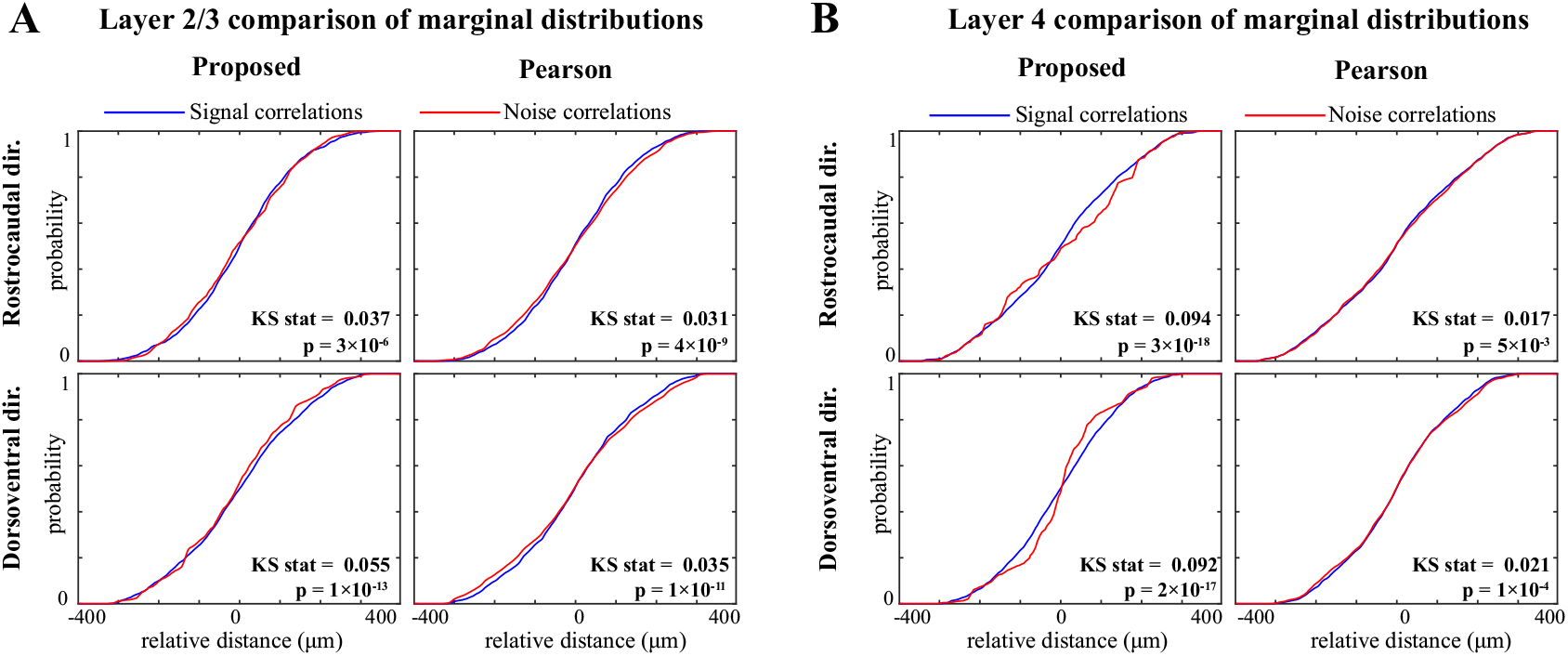
Comparison of marginal distributions of signal and noise correlations. A) Cumulative marginal probability distributions of signal (blue) and noise (red) correlations along the rostrocaudal (top) and dorsoventral (bottom) directions, as estimated by the proposed method (left) and Pearson correlations from two-photon data (right), in layer 2/3 neurons. The Kolmogorov-Smirnov (KS) test statistic along with the corresponding p-values are indicated as insets in each panel. Panel B shows the results for layer 4 in the same organization as panel A. These results show that along both directions and in both layers, the signal correlation distributions are significantly different from the corresponding noise correlation distributions, consistently for both methods. However, the KS statistics (i.e., effect sizes) for the proposed estimate are remarkably larger than those obtained from the Pearson estimates.

**Figure 7-Figure supplement 2.**
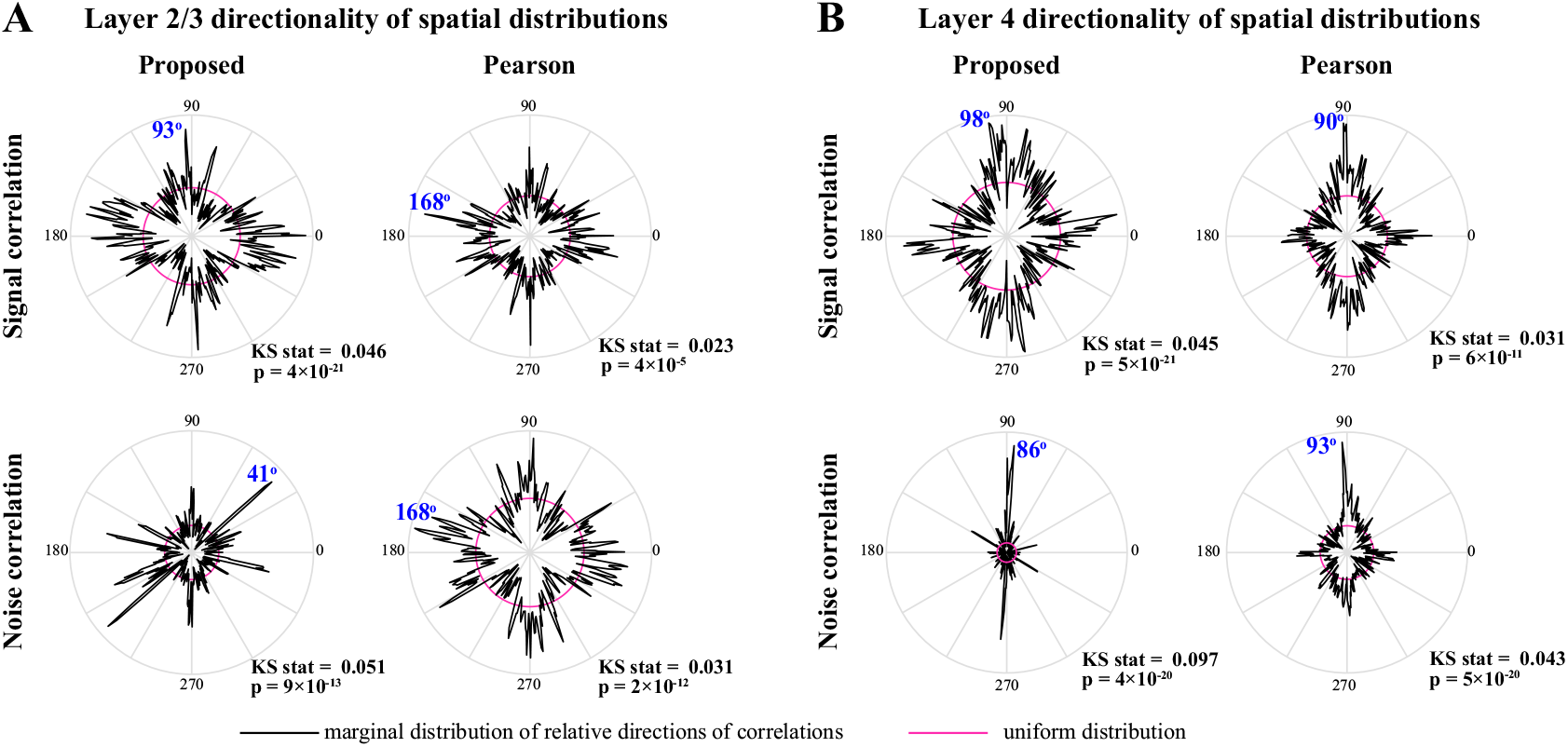
Polar plots of the angular marginal distributions of correlations. A) Polar histograms indicating the distribution of signal (top) and noise (bottom) correlations as a function of relative angle (in the dorsoventral-rostrocaudal coordinate system) between pairs of neurons in layer 2/3, as estimated by the proposed method (left) and Pearson correlations from two-photon data (right). The KS test statistic comparing each polar distribution with a uniform distribution (shown in magenta), along with the corresponding p-values are indicated below each polar plot. The mode of each probability distribution is also indicated in blue fonts. Panel B shows the results for layer 4 in the same organization as panel A. All distributions are significantly nonuniform, and particularly indicate a rostrocaudal directionality in layer 4 (as indicated by the mode angles in panel B).

